# Visualization of Incrementally Learned Projection Trajectories for Longitudinal Data

**DOI:** 10.1101/2022.11.25.515889

**Authors:** Tamasha Malepathirana, Damith Senanayake, Vini Gautam, Martin Engel, Rachelle Balez, Michael D. Lovelace, Gayathri Sundaram, Benjamin Heng, Sharron Chow, Chris Marquis, Gilles Guillemin, Bruce Brew, Chennupati Jagadish, Lezanne Ooi, Saman Halgamuge

## Abstract

Longitudinal studies that continuously generate data enable the capture of temporal variations in experimentally observed parameters, facilitating the interpretation of results in a time-aware manner. We propose IL-VIS (Incrementally Learned Visualizer), a new machine learning pipeline that incrementally learns and visualizes a progression trajectory representing the longitudinal changes in longitudinal studies. At each sampling time point in an experiment, IL-VIS generates a snapshot of the longitudinal process on the data observed thus far, a new feature that is beyond the reach of classical static models. We first verify the utility and correctness of IL-VIS using simulated data, for which the true progression trajectories are known. We find that it accurately captures and visualizes the trends and (dis)similarities between high-dimensional progression trajectories. We then apply IL-VIS to longitudinal Multi-Electrode Array data from brain cortical organoids when exposed to different levels of Quinolinic Acid, a metabolite contributing to many neuroinflammatory diseases including Alzheimer’s disease, and its blocking antibody. We uncover valuable insights into the organoids’ electrophysiological maturation and response patterns over time under these conditions.

## Introduction

Longitudinal studies offer an important perspective on evolving biological systems in domains such as microbiology, viral epidemiology, and developmental etiology. By visualizing the data generated from such evolving systems at different time intervals, analysts can piece together a more complete picture of the aspects of interest and their journey through time. Owing to the typically large number of variables considered in these studies, dimensionality reduction often plays a key role in data visualization. The fundamental concept of dimensionality reduction is to transform a distance matrix defined on a high-dimensional dataset into a low-dimensional space. A multitude of dimensionality reduction techniques exists for visualization, from methods based on statistical and linear methods (e.g., PCA^1^ and MDS^2^) to that based on topological analysis (e.g., t-SNE^3^ and UMAP^4^).

However, frequently used visualization methods such as t-SNE and UMAP are non-parametric, meaning that they do not retain a parametric model for future use with batches of subsequently obtained data which is a crucial necessity in longitudinal studies. Consequently, the visualizations generated by these methods at different sampling time points can change drastically with each other when new data is incrementally added (Fig. 2B). This volatility to incremental data undermines the reliability of insights and interpretations derived from these methods. Our previous work, Self-Organizing Nebulous Growths (SONG)^5^ was proposed with the need for a parametric visualization method in mind. SONG is capable of integrating multiple batches of new data while generating progressive/evolving visualizations, without unduly distorting the already inferred knowledge. Therefore, we extend the ability of parametric dimensionality reduction methods such as SONG to derive insights into longitudinal studies.

We propose IL-VIS (Incrementally Learned Visualizer), a machine-learning pipeline that aims to visualize longitudinal data obtained from cortical organoids, providing insights into the progression of their electrophysiological properties as shown in Fig. 1. We hypothesize that the progression of the electrophysiological properties of an organoid can be represented as a trajectory (*T*^*H*^) of the high-dimensional Multi-Electrode-Array (MEA) data. We aim to obtain a low-dimensional (2D) representation (*T*^*L*^) of *T*^*H*^ while keeping the longitudinal relationships intact to study the temporal changes of the electrophysiological properties over the experimented period. A good visualization method should be able to project *T*^*H*^ to *T*^*L*^ as accurately as possible while implicitly capturing the time axis as shown in Fig. 1C. IL-VIS employs an incremental learning approach, wherein it learns about MEA data belonging to one sampling timepoint of an organoid at-a-time (Fig. 1). This incremental process enables IL-VIS to generate visualizations of ongoing experiments with the existing data and generate improved visualizations when new data of the same experiment is available.

**Figure 1.**
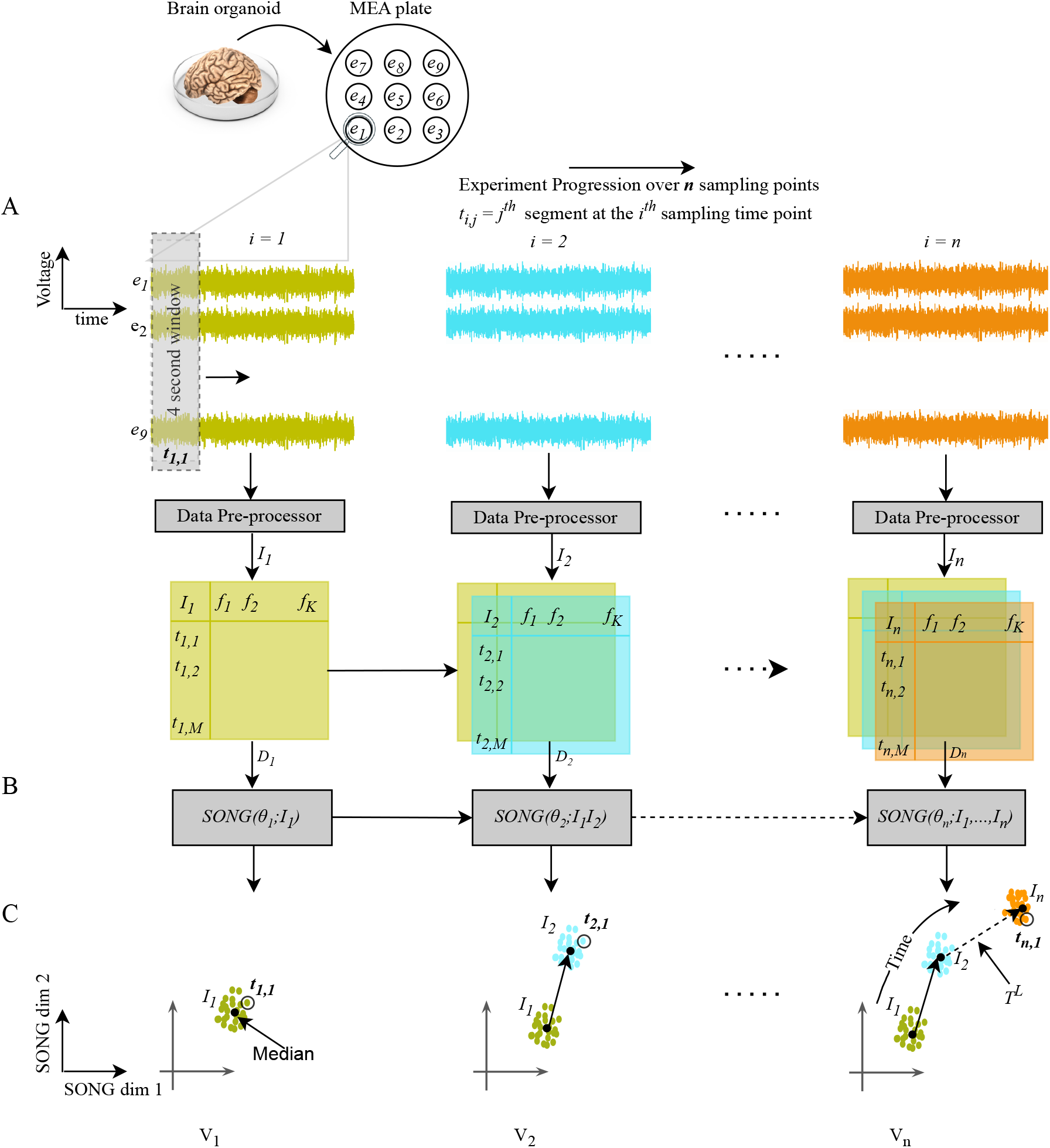
Schematic representation of “IL-VIS” using (A) MEA signals from *n* sampling timepoints, (B) the incrementally learned dimensionality reduction model, and (C) 2D partial visualizations generated to cover the progression of electrophysiological properties of the organoid thus far. (A) The activity of the brain organoid is recorded by 9 electrodes (*e*_1_ - *e*_9_) of an MEA plate in *n* sampling timepoints. These recordings are segmented into 4-second intervals. Each segment, denoted as *t*_*i*, *j*_ (*j*^*th*^ signal segment of the *i*^*th*^ sampling timepoint), is pre-processed using FFT (Methods). The resulting frequency components for each segment are represented as the segment’s features (*f*_1_, …, *f*_*k*_). *I*_*i*_ is a collection of *M* pre-processed segments recorded at the *i*^*th*^ sampling timepoint (*I*_*i*_ = {*t*_*i*, *j*_ for *jε{*1,.., *M*}}). The *n* increments corresponding to the *n* timepoints, are shown in different colors that are consistent throughout the diagram. (B) The dimensionality reduction model is trained for *n* sessions, once for each sampling timepoint in the biological experiment. At session *i*, the model parameters *θ*_*i*_ are learned by updating the parameters from the previous session (*θ*_*i−*1_) using 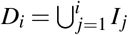, and a partial visualization *V*_*i*_ is generated. (C) Each data point in a partial visualization corresponds to a compact representation of a 4-second segment. The directed arrows map the medians of the increments learned so far in chronological order to visualize the trajectory representing the progression of the organoid’s electrophysiological properties up to that point in time. Note that none of the axes corresponds to time but the time axis is implicit in the visualized trajectory.

**Figure 2.**
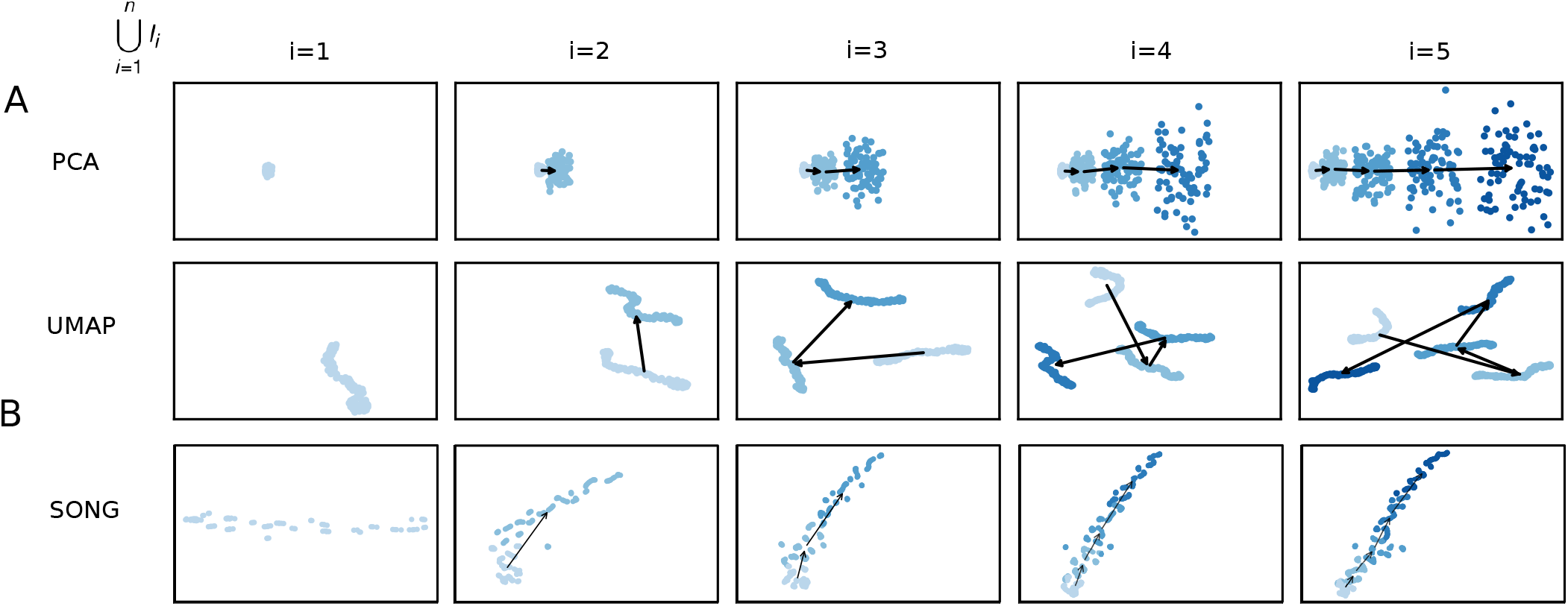
2D visualizations obtained by the pipeline when modeling 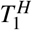 using (A) PCA, (B) UMAP and (C) SONG as the dimensionality reduction technique. 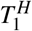 was defined on a pseudo-temporal parameter *t* (Equation 2 in Methods). *ϕ* (*t*) is noise sampled from a Gaussian distribution (see Methods). *λ* is the percentage of noise added which is 20% for this experiment. Five increments of data points are formed (*n* = 5) for each trajectory with no sampling gaps (*δ* = 0) in between thus each increment contains 100 data points (see Methods). *N* corresponds to the number of data points used in each visualization. Each row contains the five visualizations generated at the five sessions where a new data increment is introduced at each session. At session *i*, all the increments thus far (i.e.,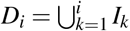) are visualized. In visualizations with multiple increments, different increments are displayed in varying intensities of the same color. The stronger the intensity, the newer the increment. By tracing the medians of the subsequent increments chronologically, the emergence of 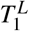 which is a 2D representation of 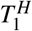 is observed. (A) PCA generated progressive visualizations but failed to preserve the non-linearity of 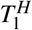 (B) UMAP preserved the non-linearity of 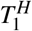 but failed to generate progressive visualizations over the sessions. (C) SONG preserved the non-linearity while generating progressive visualizations over the sessions.

First, we validate IL-VIS on simulated data to demonstrate that it accurately projects the original high-dimensional trajectories onto 2D visualizations while capturing their trends and (dis)similarities in progressions. Furthermore, we show that IL-VIS is robust to varying degrees of 1) noise in data and 2) gaps between the sampling time points. Second, we apply IL-VIS to data from experimental cortical organoids derived from Alzheimer’s Disease (AD) patients and healthy individuals. The generated visualizations are subsequently used to gain insights into the maturation of organoids when exposed to “Quinolinic Acid (QUIN)” - a metabolite known to play a role in many neuroinflammatory diseases including AD^6^ and its blocking antibody: “anti-QUIN monoclonal antibody (anti-QUIN)”^7^. We observe that the progression of the electrophysiological properties corresponding to the organoids exposed to anti-QUIN exclusively shows a distinct behavior compared to organoids exposed to other treatments, irrespective of the AD/healthy status. Furthermore, by comparing the progression of organoids treated with varying levels of QUIN and anti-QUIN, we find that anti-QUIN can sequester endogenous and exogenous QUIN, offering valuable insights into its therapeutic effects.

## Results

### Overview of IL-VIS

IL-VIS (Fig. 1) incrementally learns a high-dimensional trajectory *T*^*H*^ representing the progression of electrophysiological properties of an organoid over *n* sampling timepoints and visualizes it in 2D.

The *m*(= 9) signals corresponding to the *m* electrodes in the MEA at each sampling timepoint *i*, where 1 ≤ *i* ≤ *n*, are divided into segments of 4 seconds each. Each segment, denoted as *t*^*i*, *j*^ for the *j*^*th*^ segment at the *i*^*th*^ sampling timepoint, undergoes preprocessing using Fast Fourier Transformation (FFT)^8^ (See Methods). The collection of pre-processed *t*^*i*, *j*^ segments ∀ *j* forms the *i*^*th*^ data increment, represented as *I*_*i*_ (illustrated in Fig. 1A). *I*_*i*_ is then reduced to 2D using the incrementally trained dimensionality reduction model based on SONG (Fig. 1B).

The model is trained for *n* sessions, once for each sampling timepoint. At session *i, I*_*i*_ is combined with the existing Data 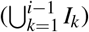, and the combined dataset 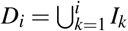 is used to train the existing model parameters *θ*_*i−*1_ to produce updated parameters *θ*_*i*_. *θ*_*i*_ is then used to generate a 2D visualization *V*_*i*_ that visualizes all the increments from *I*_1_ to *I*_*i*_ (Fig. 1C). The median of each *I*_*i*_ in *V*_*i*_ is traced from 1 to *i* in order to obtain *T*^*L*^ that visualizes the progression of the organoid’s electrophysiological properties over time. If *n* is sufficiently high, we would expect the consecutive and continuous data increments from an organoid to form a gradual progression and appear seamlessly connected, generating a continuous visualization. In the absence of such a continuous stream of data, a good visualization method should identify the trajectory progression even using discrete increments.

When visualizing the electrophysiological progression of a single organoid, the direction of the trajectory cannot be compared to another independently visualized organoid. Therefore, we extend IL-VIS to jointly model multiple organoids, allowing us to examine the directions of trajectories compared to each other and identify (dis)similarities in the electrophysiological progression of the corresponding organoids (see Figs. 6A-D, explained later).

In subsequent sections, we report two sets of results using 1) simulated data and 2) experimental MEA data obtained from cortical organoids.

### Simulated data

Several studies have observed a gradual and non-linear increase in the electrical activity of brain organoids as they mature, characterized by parameters such as spike frequency, bursts, and synchrony^9,10^. This increase in activity is indicative of the development of more complex and mature synaptic connections among neurons and stronger electrical transmission within the organoids^11^. We verify IL-VIS’s ability to visualize this gradual but non-linear progression of electrophysiological properties as captured in high-dimensional data over time, in a 2D representation. However, it is possible that the actual progression of the electrophysiological properties may not show a gradual progression. This could be due to various factors such as developmental pathology, the influence of a disease, or administered therapeutic stimuli. After confirming that IL-VIS can accurately capture the expected gradual increase trend in healthy organoids, we can use IL-VIS to identify deviations from these trends resulting from the factors described above. Next, we validate IL-VIS’s capability to capture the (dis)similarities between multiple progression trajectories. This feature allows for visualization of the effect of different treatments on the electrophysiological progression of multiple organoids relative to each other.

We generate simulated data for which the corresponding high-dimensional trajectory *T*^*H*^ of the visualized trajectory *T*^*L*^ is known. Specifically, we formulate a set of non-linear high-dimensional trajectories {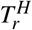 where *rε*{1,…, 4}} that gradually progress with time. The details related to the formation of *I*_*i*_, 1 ≤ *i* ≤ *n* in each 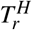 can be found in Methods.

#### IL-VIS generates evolving visualizations capturing the underlying trends

Fig. 2 shows the visualizations obtained using PCA and UMAP in place of SONG in IL-VIS in which PCA fails to capture the non-linearity present in data, while UMAP fails to generate evolving visualizations across the sessions. Additional experiments with Independent Component Analysis (ICA), Multi-Dimensional Scaling (MDS), and t-distributed Stochastic Nonlinear Embedding (t-SNE) are provided in the supplementary Fig. 2. These experiments demonstrate that they either fail to preserve the non-linearity, generate evolving visualizations (by preserving the orientation of the trajectories), or both. In contrast, by using SONG, as shown in Figs. 3 and 4, visualizations successfully capture the gradually increasing trend and the non-linearity of the high-dimensional trajectories (see Supplementary Fig. 1 for pairwise Euclidean and geodesic distance heatmaps between the increments in 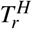and 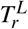). Significantly, IL-VIS also generates evolving visualizations, preserving the relative structure between the increments or the orientation across incremental visualizations, confirming that IL-VIS could be used for gaining insights from ongoing experiments. This ability stems from two key factors. Firstly, SONG operates as a parametric method, and secondly, IL-VIS preserves and reuses these parameters throughout iterations. To elaborate further, we conducted an additional experiment in which we reset the parameters at the beginning of each session, intentionally excluding the carryover of parameter values from the previous session. The resulting visualizations are shown in Supplementary Fig. 3. Notably, across sessions, the orientation of the trajectories is not preserved when the parameters are not carried forward. While it is conceivable to manually reorient the trajectories of each visualization, this approach may be impractical for intricate trajectories, highlighting a limitation of this approach. This underscores the significance of parameter continuity in preserving the meaningful evolution of visual representations over time. Additionally, we verify the manifold preservation quality of the SONG embeddings in Supplementary Fig. 4, and the method’s capability to cater to an even higher dimensional space (up to 10000) in Supplementary Fig. 5.

**Figure 3.**
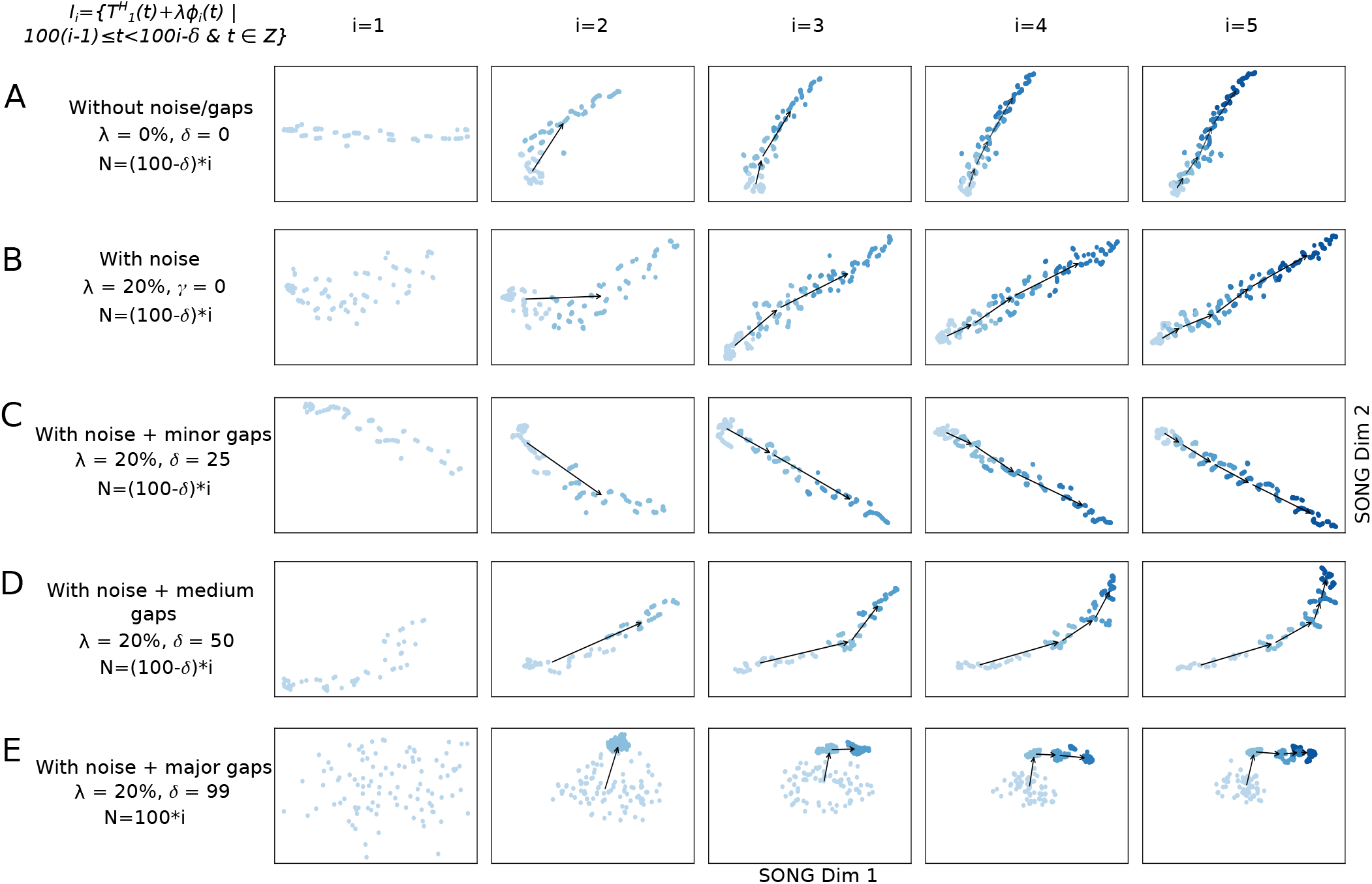
2D visualizations obtained by IL-VIS showed its capability in generating evolving visualizations capturing the gradually increasing and non-linear trends of a simulated 100-dimensional non-linear trajectory 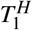 (A) without noise/gaps, (B) with noise, (C) with noise and minor sampling gaps between increments, (D) with noise and medium sampling gaps between increments, and (E) with noise and major sampling gaps between increments. 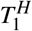 was defined on a pseudo-temporal parameter *t* (Equation 2). *ϕ*_*i*_(*t*) is noise sampled from a Gaussian distribution (Methods). *λ* is the percentage of noise added: 0% for (A) and 20% for (B)-(E). For each experiment, five data increments were formed (*n* = 5). The number of data points in each increment was determined based on *δ* and was 100, 100, 75, 50, and 100 for experiments A-E respectively (see Methods). *N* corresponds to the number of data points used in each visualization. Each row contains the five visualizations generated at the five sessions where a new data increment is introduced to IL-VIS at each session. At session *i*, all the increments thus far (i.e., 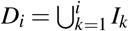) are visualized. In visualizations with multiple increments, different increments are displayed in varying intensities of the same color. The stronger the intensity, the newer the increment. By tracing the medians of the subsequent increments chronologically, the emergence of 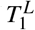 which is a 2D representation of 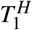 is observed. Each consecutive visualization has maintained the structure across the previous visualization allowing us to relate between the visualizations. Furthermore, the progressiveness of the gradually increasing trajectory is captured by placing each new increment further away from the previous increments (Supplementary Fig.1). IL-VIS shows robustness to noise (B) and minor to major gaps (C)-(E).

**Figure 4.**
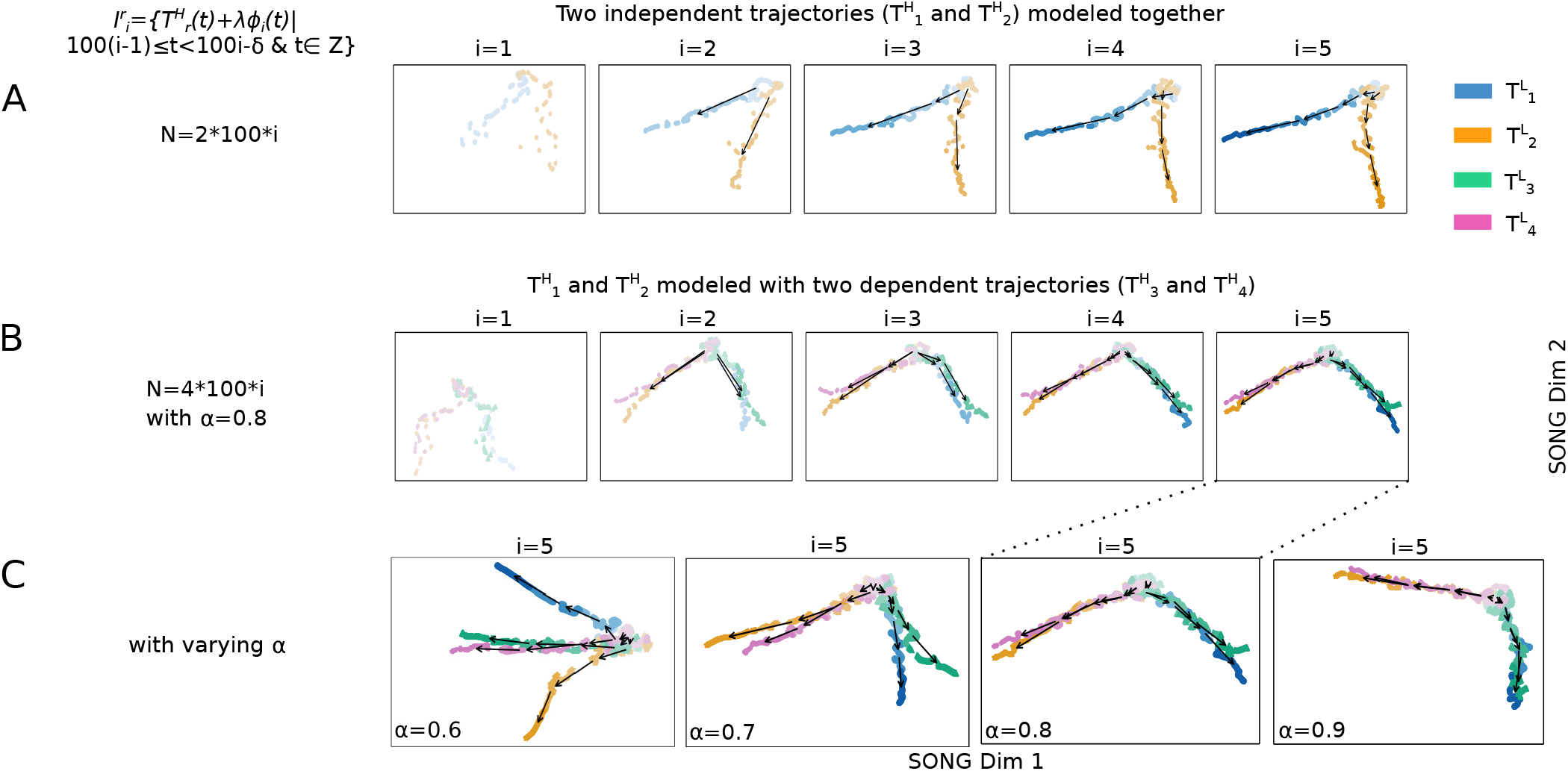
2D visualizations obtained by IL-VIS showed its capability in capturing the (dis)similarities in the progressions of (A) two trajectories: 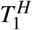 and 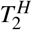 and (B) four trajectories: 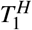 and 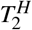 together with 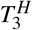 and 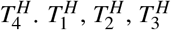, and 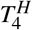 are 100-dimensional non-linear trajectories. 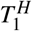 and 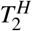 are independent (principle trajectories) whereas 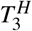 and 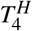 are dependent (secondary trajectories) on 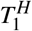 and 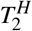. Specifically, 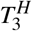 and 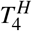 are created as compositions of 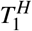 and 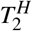 where a similarity factor (*α*) is used to determine how closely a secondary trajectory is to a principal trajectory (see Equation 3). *ϕ*_*i*_(*t*) is noise sampled from a Gaussian distribution and *λ* is the percentage of noise added which is 0% for this experiment (Methods). Five increments of data points are formed (*n* = 5) for each trajectory with no sampling gaps (*δ* = 0) in between thus each increment contains 100 data points (Methods). 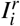 is the *i*^*th*^ data increment of 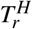. *N* corresponds to the total number of data points used in each visualization. Each experiment contains five visualizations generated at the five sessions. At each session *i*, the *i* data increment of all trajectories that are modeled together is introduced to the pipeline. Each trajectory is shown in a different color. In visualizations with multiple increments, different increments are displayed in varying intensities of the same color. The stronger the intensity, the newer the increment. By tracing the medians of the subsequent increments chronologically, the emergence of 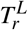 which is a 2D representation of 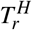 is observed. (A) 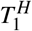 and 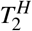 originate at the same position and diverge from each other as the increments are added which is consistent with the definition of the trajectories (B) when *α* = 0.8, 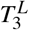 is visualized closer to 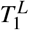 than to 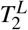, and 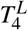 is visualized closer to 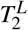 than to 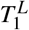 capturing the relative similarity between the principle and secondary trajectories (C) Visualizations generated at the last session (*i* = 5) in four independent experiments with four different *α* values. The *α* was varied from 0.6 to 0.9 to observe the movement of 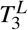 and 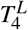 in reference to 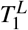 and 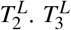 and 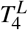 are visualized closer to 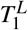 and 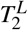 respectively as the *α* is increased. Altogether, visualizations show IL-VIS’s capability in capturing the (dis)similarities in trajectory progressions.

**Figure 5.**
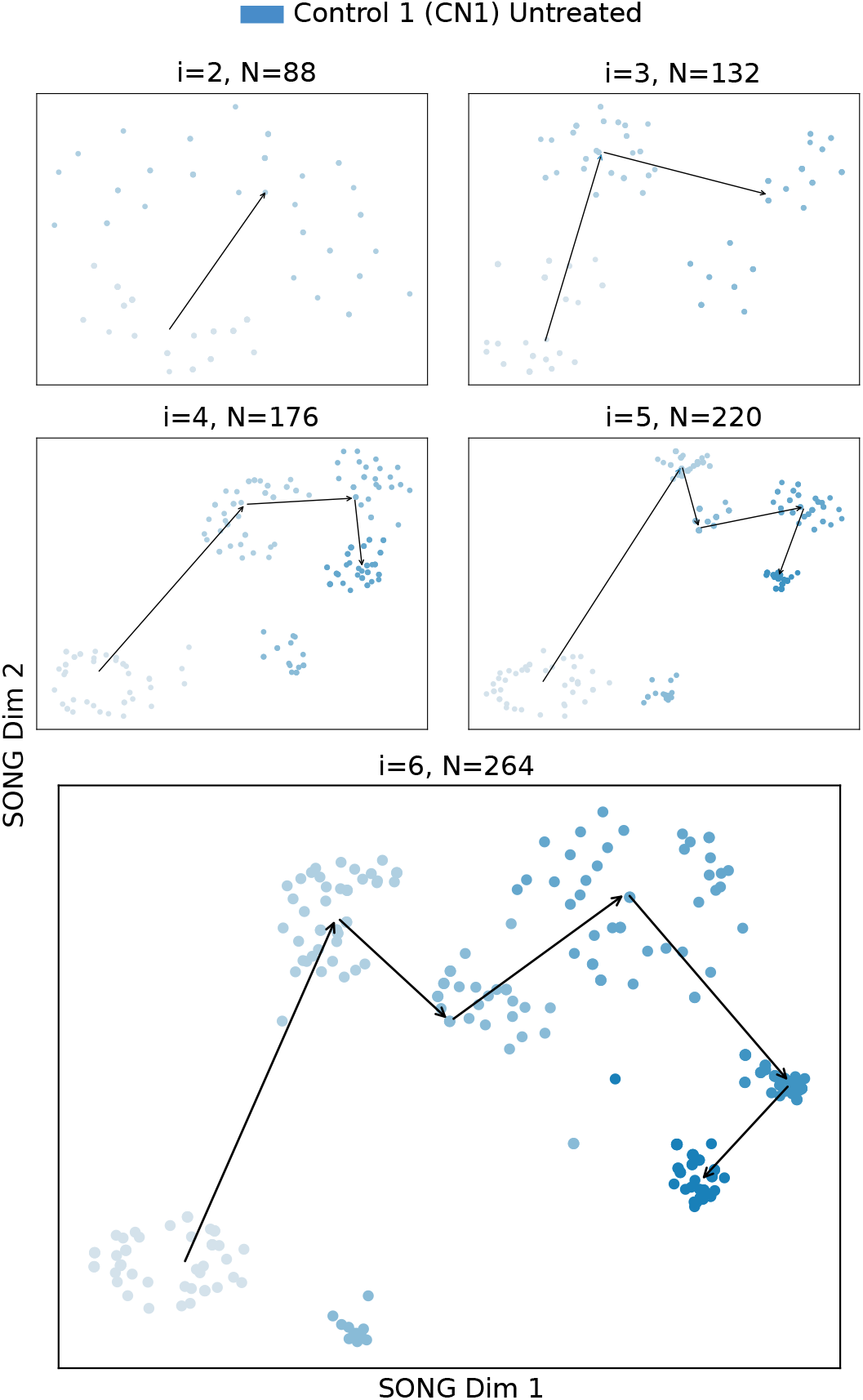
2D visualizations obtained by IL-VIS for the Untreated CN1 organoid when modeled independently. *I*_1_ to *I*_6_ are the six increments corresponding to the six different time points at which the recordings were taken (Table 2). At session *i*, all the increments thus far (i.e.,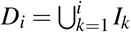) are used for training the model. *N* corresponds to the number of data points used in each visualization where each point corresponds to a compact representation of a 4-second recording obtained from the 9 electrodes in the respective MEA plate. The 2D visualizations generated from the 2^*nd*^ session (*i* = 2) onwards are shown in the sub-figures. The color intensity represents the organoid’s maturity. In earlier increments, the color is lighter while in later increments the color is darker. The medians of the subsequent increments are traced chronologically to observe the trajectory followed by the organoid’s electrophysiological properties.

**Figure 6.**
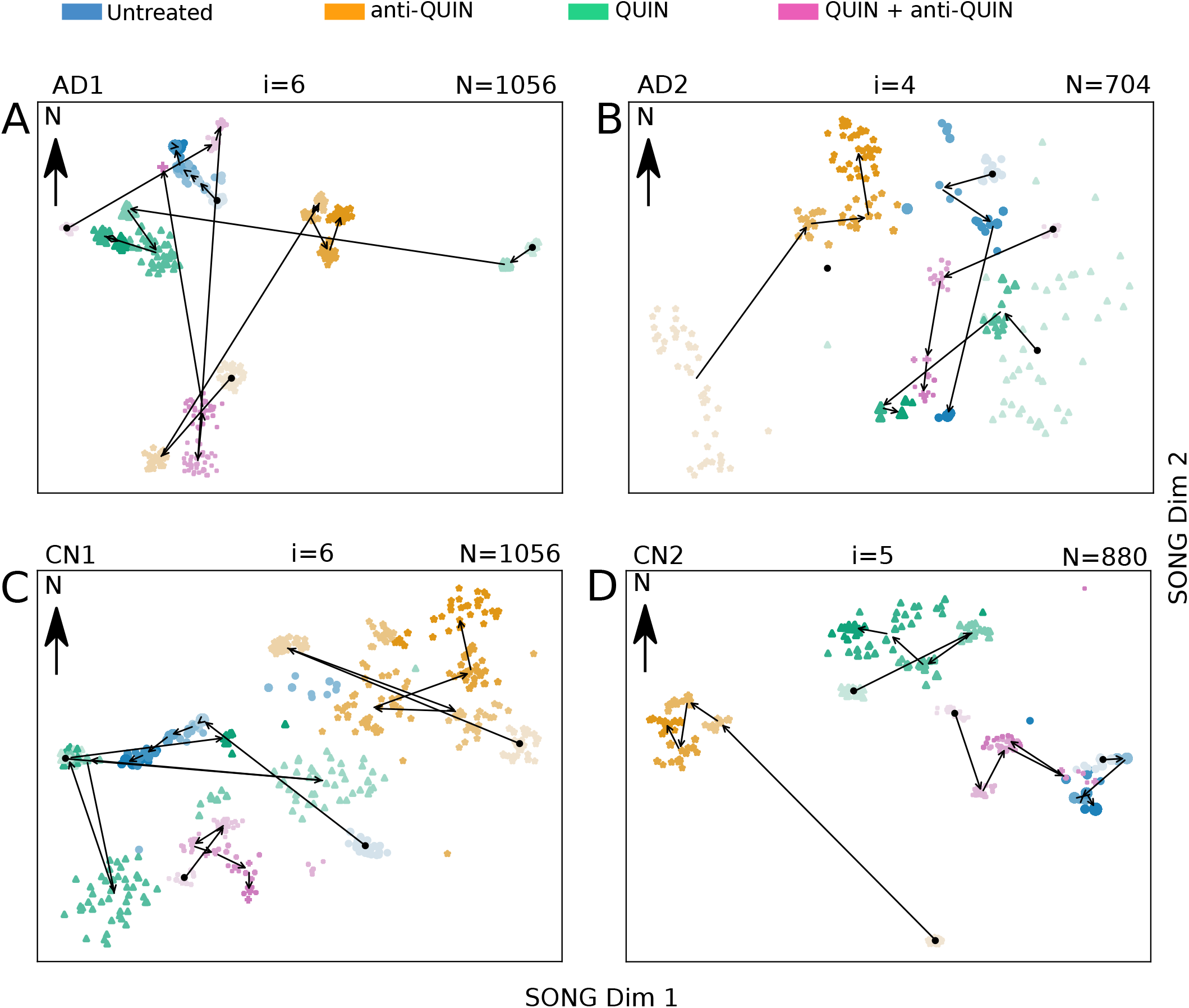
The 2D visualizations obtained by IL-VIS for each cell line at the end of the three months treatment period. The MEA data from the four organoids of the same cell line, treated with Untreated, QUIN, anti-QUIN, and QUIN+anti-QUIN, are modeled together. The data point representation is the same as Fig. 5. The visualizations show a unique progression in AD/CN organoids treated with anti-QUIN alone, compared to the progression of Untreated, QUIN-treated, or QUIN+anti-QUIN-treated organoids. (A) The anti-QUIN-treated trajectory originates at a considerable distance and moves in a different direction from the trajectories corresponding to the other three treatments. In contrast, both Untreated and QUIN+anti-QUIN treated trajectories initiate and converge in nearby locations to each other. Although the QUIN-treated trajectory initiates at a considerable distance, it converges to a point closer to the Untreated and QUIN+anti-QUIN-treated trajectories. (B) anti-QUIN-treated trajectory originates from the bottom left corner of the figure at a considerable distance from the three trajectories corresponding to the other three treatments which are originating at the right side of the figure. anti-QUIN treated trajectory propagates in the opposite direction (SW to NE and then towards N) than the other trajectories and converges in a distant location. The trajectories corresponding to the other three organoids converge closer to each other. (C) The anti-QUIN treated organoid is originating at a considerable distance from the other three organoids and moves in a different direction (top right). In contrast, the other three organoids are located more toward the bottom left corner of the figure. (D) The anti-QUIN treated organoid is originating at a considerable distance from all the other three organoids and moves towards the top left of the figure. In contrast, the other three organoids are located more toward the top right corner of the figure.

#### IL-VIS is robust to noise and gaps in the data

To replicate random noise arising from biological and non-biological variations in real data, we add random Gaussian noise (20%) to each data increment in 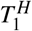(see Methods) and visualized in Figs. 3B-E. Despite the presence of this added noise, 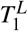successfully preserves the trend and non-linearity of 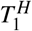 (Figs. 3B-E). However, as the noise level increases from 0 to 20%, there’s a corresponding decrease in the smoothness of 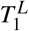 (Supplementary Fig. 6).

Although the processes underlying the biological experiments are continuous, the recorded features are discrete and the sampling time points may have considerable gaps between them. To replicate this, we introduce minor, medium, and major gaps between the increments in 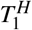 (Methods) and visualize them in Figs. 3C-E. Despite such gaps, 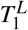 captures a gradually increasing and non-linear path along which 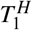 progresses, suggesting that IL-VIS can handle datasets with considerable gaps and noise. When major gaps are introduced (where *δ* = 99), five different clusters are visualized each corresponding to an increment instead of a continuous progression (Fig. 3E). This agreed with our expectations because IL-VIS had very limited information about continuity.

#### IL-VIS captures (dis)similarities between the trajectory progressions

The progression of electrophysiological properties of multiple organoids with different exposures may correspond to different trajectories in the high-dimensional space. Capturing the (dis)similarities between these trajectories and visualizing them as they evolve in a shared space would offer insights into the (dis)similarities in their electrophysiological progressions. For instance, a comparison between the electrophysiological maturation of organoids exposed and not exposed to a particular treatment would assist in understanding the treatment’s effect. We validate this capability of IL-VIS as below.

First, we jointly model and plot two independent randomly-generated non-linear trajectories (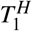, and 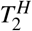) designed to originate at the same position while diverging at higher *t* values in Fig. 4A. As the new increments are added, the corresponding low dimensional trajectories 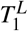 and 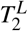 exhibit an increasing difference, highlighting their growing dissimilarity in later increments. Next, we extend the experiment by incorporating two secondary trajectories 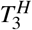 and 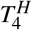, along with the principal trajectories (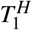 and 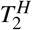). 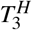 and 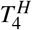 are created as combinations of 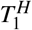 and 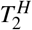, based on a similarity factor *α* (Methods). Specifically, when *α* = 1, 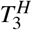 coincides with 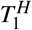, and 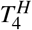 coincides with 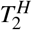. Conversely, when *α* = 0, 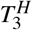 coincides with 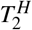, and 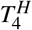 coincides with 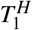. For 0 < *α* < 1, 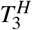 and 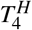 would be exhibiting intermediate characteristics between 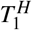 and 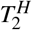. along the trajectory continuum. The resulting 2D visualizations in Figs. 4B and 4C showcase these distinctive traits in the low-dimensional trajectories. Notably, when *α >* 0.5, 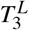is visualized closer to 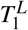 than 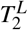, and conversely, 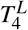is visualized closer to 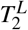 than 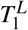 (Supplementary Table 1). Subsequently, in Fig. 4C, as *α* is incremented from 0.6 to 0.9, the visualizations of 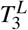 and 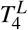 gradually approach 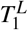 and 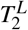, respectively. Furthermore, within trajectory pairs such as 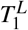 and 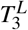 sharing similarities, i.e., when *α* > 0.5, the trajectories display increased separation in later increments compared to earlier increments, effectively capturing subtle but important dissimilarities. The experiments described above combined with these results, validate IL-VIS’s ability to generate evolving visualizations that preserve the high-dimensional trajectories’ similarity relationships.

### Cortical organoids data

After validating IL-VIS on simulated data, IL-VIS is then applied to MEA data obtained from cortical organoids. MEA data has been previously used to assess alterations in neuronal excitability, e.g., mediated by neuroinflammation^12^. QUIN is an N-methyl-D-aspartate (NMDA) agonist generally synthesized and released during neuroinflammation. QUIN’s over-activation of NMDA receptors leads to neuronal excitotoxicity, thus, QUIN has been implicated in a range of diseases in which neurodegeneration is a key component, including AD, multiple sclerosis, and stroke^13–16^. QUIN has also recently been shown to reduce both neurite outgrowth and the number of synaptic spines in culture^17^. QUIN antibodies can provide neuroprotection^7^ by preventing QUIN binding to NMDA receptors or its uptake offering therapeutic potential for these diseases. In a previous study^7^, we showed that as little as 30 minutes of exposure to the same monoclonal anti-QUIN used in this study could protect oligodendrocyte cell line monolayers from cell death induced by QUIN. QUIN excitotoxicity and metabolism are very much associated with the aging process^18,19^, thus, the long-term effect of anti-QUIN to mitigate the effects of QUIN should be studied. Therefore, we analyze both the acute and chronic effects of exogenous anti-QUIN on both endogenous and exogenous QUIN using IL-VIS.

Our dataset (see Methods) contains MEA data from cortical organoids derived using four cell lines: two AD patients (AD1 and AD2) and two healthy controls (CN1 and CN2). Four organoids are derived from each cell line and exposed to one of the following four different treatments 1) Untreated, 2) QUIN added, 3) anti-QUIN added, and 4) both QUIN and anti-QUIN added (QUIN + anti-QUIN). These 16 organoids from 4 cell lines are used to test the ability of anti-QUIN to sequester, thus reducing the deleterious effects of QUIN. QUIN is also produced by the organoids endogenously and released into the media, where it was quantified at 16-30 nM (Supplementary, Fig.8). The untreated organoids are used as controls for treatments 2), 3), and 4), and to establish the normal maturation profile reflected in the plots discussed later in Fig. 6. In treatment 2), organoids are exposed to exogenous QUIN at 100 nM, i.e., 3-5 times endogenous levels but lower than the circulating levels in blood^20^. This concentration is chosen to ensure physiological relevance to test the effect of increased QUIN on organoids. In treatment 3), organoids are exposed to anti-QUIN alone to determine anti-QUIN’s ability to sequester endogenous levels of QUIN. In treatment 4), organoids are exposed to both anti-QUIN and exogenous QUIN to assess anti-QUIN’s capability to sequester higher levels of QUIN. In treatments 3) and 4), anti-QUIN is added at 10 *µ*g*/*mL concentration to reduce extracellular QUIN concentrations and prevent QUIN uptake and binding to NMDA receptors.

#### IL-VIS identifies the progression of the electrophysiological properties of an organoid

The progression of the electrophysiological properties of the untreated organoids, originally represented as high-dimensional trajectories in MEA data 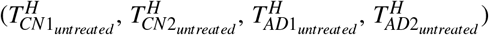 are transformed into 2D trajectories 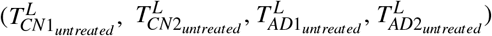 using IL-VIS (Fig. 5 and Supplementary Figs. 9-11). In contrast to a conventional spike count analysis (Supplementary Fig.12), the generated visualizations offer a new perspective on the changes in an organoid’s electrophysiological state over time compared to its previous states. For instance, in Fig. 5, the 2^*nd*^, 3^*rd*^, and 4^*th*^ increments of 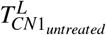 appear to move away from the 1^*st*^. However, the 5^*th*^ and 6^*th*^ increments reverse the direction and move back towards the initial state, indicating that the organoid’s final electrophysiological states are becoming more similar to the initial state. These visualizations allow us to visualize the electrophysiological progression in individual organoids.

#### IL-VIS visualizes the progression of multiple organoids in the same visualization space

The spike frequency variation over the experimental period is depicted in Supplementary Fig. 12. However, it only shows the independent temporal variation and does not allow for a direct comparison of the effects between different treatments on organoids. We use IL-VIS to visualize the electrophysiological progression of the four organoids derived from the same cell line but exposed to different treatments in a shared space, enabling their comparison. Fig. 6 displays these visualizations obtained for each cell line at the end of a three-month treatment period.

The proximity between low-dimensional trajectories indicates the similarity in the electrophysiological progression of the corresponding organoids during their maturation. For instance, the increments in 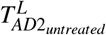 and 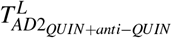 (Fig. 6B) are visualized closely, signifying the similar progressions of the untreated and QUIN+anti-QUIN-treated organoids derived from the AD2 cell line. We use an Euclidean distance-based metric (Methods) to quantitatively assess the proximity between trajectories, which provides a measure of the similarity in electrophysiological progression among the organoids.

#### The anti-QUIN-treated organoid trajectory deviates from the other three conditions in all four cell lines

Fig. 6 illustrates that in each cell line, the trajectory corresponding to the anti-QUIN-treated organoid originates and progresses at a considerable distance from the trajectories corresponding to the other three conditions (Untreated, QUIN, and QUIN+anti-QUIN). Additionally, the trajectory of the anti-QUIN-treated organoid progresses in a distinctly different direction compared to the trajectory of the QUIN-treated organoid. For instance, in Fig.6A, 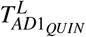 progresses from E to W while the 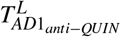 progresses from SW to NE. Similarly, in Fig. 6B, 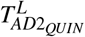 (NE to SW) and 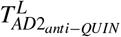 (SW to NE) are progressed in parallel but in opposite directions. These findings suggest that anti-QUIN has a unique impact on electrophysiological activity compared to the other treatments.

#### Possible anti-QUIN sequestration of QUIN

The distances measured to trajectories of Untreated, QUIN, and QUIN+anti-QUIN-treated organoids by taking the trajectories of anti-QUIN-treated organoids as reference are ranked in increasing order in Table 1. For three of the four cell lines: AD1, AD2, and CN1, the treatments were ranked as follows: (1) Untreated, (2) QUIN+anti-QUIN, and (3) QUIN. As expected, the trajectory of the QUIN-treated organoid is the least similar (ranked 3^*rd*^) to the trajectory of the anti-QUIN-treated organoid. The trajectory of the untreated is the closest (ranked 1^*st*^) while the trajectory of the QUIN+anti-QUIN-treated is the second closest (ranked 2^*nd*^). By attributing this closeness to the level of the effective concentration of QUIN bathing the organoids, we conjecture that the order of effective concentration of QUIN in organoids is increasing in the specific order: anti-QUIN-treated < Untreated < QUIN+anti-QUIN treated < QUIN-treated, suggesting that,

**Table 1.**
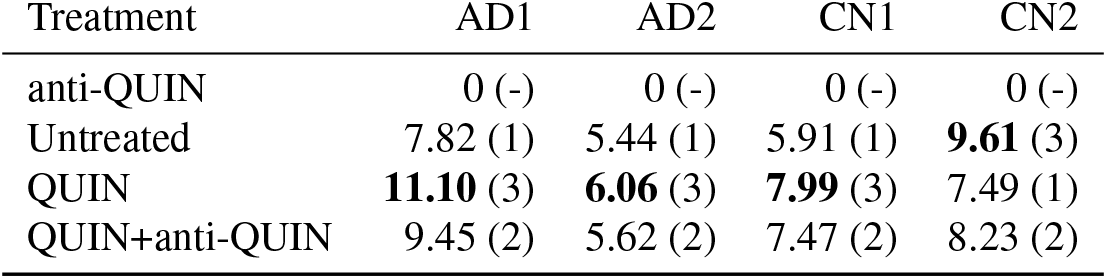
The distance measured to untreated, QUIN, and QUIN+anti-QUIN-treated trajectories by taking the anti-QUIN treated trajectory as reference. Distance rank is shown in brackets.

- The concentration levels of anti-QUIN in anti-QUIN-treated are sufficient to sequester low levels of endogenous QUIN found in Untreated organoids.
- Although not being able to fully neutralize the effect of QUIN, the concentration levels of anti-QUIN can sequester some exogenous QUIN found in QUIN+anti-QUIN-treated organoids.

For the remaining cell line (CN2), further investigation is needed to confirm the reason for the trajectory of the anti-QUIN-treated organoid being visualized closer to the trajectory of QUIN-treated than to the other two treatments and in particular to the Untreated organoid. One possible explanation for this would be that the endogenous QUIN in the Untreated CN2 is much higher after nine months than in the other three cell lines (Supplementary Fig.8). However, for CN2, the placement of the trajectories of Untreated, QUIN-treated, and QUIN+anti-QUIN-treated organoids in Fig.6 is consistent with the other three cell lines, i.e., the trajectory of QUIN-treated is closer to the trajectory of the QUIN+anti-QUIN-treated than to the trajectory of the Untreated organoid.

## Discussion

We proposed IL-VIS, a machine-learning pipeline for visualizing high-dimensional trajectories in longitudinal MEA data in a 2D space.

IL-VIS has four key advantages that are important in the fields of neurodevelopmental, pharmacological, and disease modeling studies and we demonstrate these advantages in our results section by using simulated and experimental data. First, by relying less on human intervention for feature extraction, we overcome several weaknesses in conventional MEA analysis that manually extracts features such as spike rates, and burst rates^21,22^. Such methods require expertise in choosing parameters and may introduce expert biases that would influence all subsequent analysis tasks as discussed in a previous work^23^. Second, IL-VIS can model the data in an ongoing experiment and progressively enhance the model with the arrival of new data. This eliminates the need for the initial set of data to offer a complete representation of the underlying processes that generate data. Third, IL-VIS generates intuitive and interpretable 2D representations capturing the changes in progressions in the electrophysiological activity of an organoid over time. Fourth, by visualizing the progressions of electrophysiological properties in multiple organoids in a shared space, IL-VIS can determine the effect of different treatments on the same cell line and compare their effects to each other. IL-VIS makes the third and fourth advantages possible by preserving the heterogeneity of responses at any given timepoint and in between different timepoints, reflecting anything from small to large variations in electrophysiological properties. Moreover, we show the pipeline’s robustness to gaps and noise in data which are frequently observed in experimental data.

Our application of IL-VIS on experimental data allows for the identification of the effect of QUIN and anti-QUIN on the electrophysiological activity of organoids. Results indicate that low levels of QUIN may induce effects on cellular health, and neurodegeneration, leading to electrophysiological alterations. Our findings are in agreement with recent evidence demonstrating decreased neurite growth and loss of synaptic spines in cultured neurons exposed to QUIN^17^. Furthermore, the significance of age-related changes in the kynurenine pathway in the brain, including QUIN accumulation, is starting to be appreciated in recent human studies of healthy and neurologically diseased patients^24,25^. Therefore, we believe our findings supporting the evidence of anti-QUIN’s capability in sequestering varying levels of QUIN are of greater importance. More studies are needed to longitudinally assess the levels of QUIN, the potential correlation with known neurodegeneration biomarkers, and the effect of potential interventions to reduce QUIN levels. IL-VIS can play a key role in modeling data from such future experimental studies.

IL-VIS’s potential can be tested in new applications, including the two scenarios outlined below. First, IL-VIS could be used to compare the effects of different treatments, such as conducting a relative analysis of any set of compounds (in our study it was limited to QUIN and anti-QUIN) compared to an untreated organoid. Second, the distance between individual increments of an organoid’s trajectory under a specific treatment can be used to gauge the variability of responses in relation to the treatment time. However, conducting such an experiment requires a careful design with the above purpose in mind and requires generating data at regular and frequent intervals. In addition, we acknowledge that comparing Alzheimer’s versus control cell lines is beyond the scope of this study. Such a comparison necessitates more replicates to accurately represent each disease and control population.

Further enhancements can be made to IL-VIS. We provide suggestions for improving both the machine-learning pipeline and the biological experimental design. For studies using MEAs with a larger number of electrodes, a 2D convolutional neural network can be integrated into Fourier-transformed features to leverage the spatial proximity of the electrodes. Additionally, a rehearsal-based strategy^26^, selectively storing and utilizing informative samples from previous increments, can be seamlessly integrated into IL-VIS to address limitations in storing and using all training examples. Furthermore, in contrast to dimensionality reduction techniques like PCA, where the axes correspond to the directions of maximum variance in the original high-dimensional space, manifold learning methods such as UMAP and SONG lack a direct mapping of their dimensions to specific features or characteristics. This highlights a unique challenge in manifold learning approaches. Given this distinction, future studies may delve into interpreting the intrinsic properties captured by the axes of these methods. In terms of biological experimental design, we recommend the following steps for more insightful trajectories. First, recording MEA data from multiple time points before introducing treatments would enable IL-VIS to learn and calibrate the variations among the organoids utilized. Second, minimizing the gaps between increments would enable IL-VIS to generate a smooth and continuous trajectory for each organoid. Third, utilizing varying concentrations of the same treatment on organoids would allow for a detailed analysis of the relationship between trajectories with high confidence.

In conclusion, IL-VIS presents a novel approach for analyzing high-dimensional, longitudinal MEA data, and offers promising potential to provide valuable insights into the electrophysiological progression of organoids under various treatments and conditions, ultimately advancing our understanding of complex biological processes.

## Methods

### Experimental Cortical Organoid-based MEA Data

All research was conducted in accordance with the requirements of the University of Wollongong Human Research Ethics Committee (13/299). The iPSCs utilized in this study have been described previously^27–29^ under HREC approval 13-299 (UOW). The fibroblasts were originally obtained from the Coriell Institute^27^ and the Centre of Healthy Brain Ageing (CHeBA) Research Bank under HREC approval HC17865 (UNSW). Coriell Institute ensures informed consent is obtained when collecting the samples. Participants from the CHeBA research bank were a part of the Sydney Memory and Ageing Study and/or were assessed in the clinic as detailed below. Participants took part in a detailed cognitive and medical assessment with the option to donate a sample. The participants were deemed able to provide informed consent based on clinical assessment and the informed consent was obtained from all the patients. A knowledgeable informant (close friend or family member) was also interviewed at this time. The assessment measures were (1) Neuropsychiatric assessments assessing dementia, mild cognitive assessment and cognitive decline; (2) Broad range of sociodemographic, lifestyle and cardiac, physical and mental health factors; (3) Blood samples and genotyping; (4) Subgroups with MRI scans (structural and functional) and for studies of falls and balance, inflammatory and metabolic markers, and prospective memory. These assessments were conducted by the clinicians at CHeBA at UNSW Sydney.

The dataset contains MEA recordings obtained from cortical organoids derived from four different cell lines: two healthy control cell lines (UOWi001-A and UOWi002-A) and two Familial AD patient cell lines (UOWi003-A and UOWi009-A). Both AD patient lines carry mutations in the PSEN1 gene that encodes Presenilin 1, A248E or S290C, as described in Maksour et al.^30^ and demonstrate alterations in neuronal excitability^30^. Human brain organoids^31^ produced from iPSCs based on the protocol by Salick et al.^32^ and described in Maksour et al.^29^ were cultured for up to 9 months. The organoids were plated after being cultured for five months, the treatments were added after 6 months from being plated (see Table 2) and the activity was recorded after a day, a week, a month, and three months. The durations between the recordings were selected with the intention to measure both the acute (one day) and chronic (one week, one month, and three months) effects of the treatments. The decision to include various time intervals stemmed from the unique nature of organoid cultures, which allow for extended periods of analysis with microelectrode arrays not fully exploited by the current literature because they focus on shorter time frames (of the order of days). In contrast, our sampling method aimed to capitalize on the extended culture periods possible for organoids. Furthermore, it is critical to note that our study does not involve direct comparisons between different cell lines. Instead, we focus on comparing organoids derived from the same cell line but treated with different treatments. Consequently, all organoids derived from a specific cell line share identical time intervals. While certain cell lines may exhibit experimentally missing time points, the absence or inclusion of these points among the various cell lines does not alter the outcomes of our analysis.

**Table 2.** MEA signal recording availability for each cell line.

| Organoid age | Treatment time (t) | CN1 | CN2 | AD1 | AD2 |
| --- | --- | --- | --- | --- | --- |
| 5 months | - | $I_1$ | - | $I_1$ | - |
| 6 months (Drugs added) | t=0 | $I_2$ | $I_1$ | $I_2$ | $I_1$ |
| 6 months & 1 Day | t=1 day | $I_3$ | $I_2$ | $I_3$ | - |
| 6 months & 1 Week | t=1 week | $I_4$ | $I_3$ | $I_4$ | $I_2$ |
| 6 months & 1 Month | t=1 month | $I_5$ | $I_4$ | $I_5$ | $I_3$ |
| 6 months & 3 Months | t=3 months | $I_6$ | $I_5$ | $I_6$ | $I_4$ |

The treatments were added after 6 months and the activity was recorded after a day, a week, a month, and three months. The table represents the MEA recording availability for each cell line and their corresponding increment ID ( $I_i$ )

The selection of the exogenous QUIN concentration of 100 nM was based on our previous research, which demonstrated chronic QUIN toxicity and subcellular changes including dendritic varicosities and microtubule alterations in neurons using a 350 nM dose^33,34^. Additionally, other studies have shown neuronal damage following exposure to only 100 nM QUIN for several weeks^35^. Considering the longer duration of our study (lasting three months after treatment), we opted to use a lower concentration to assess the effects of physiologically elevated levels of QUIN, while likely retaining a substantial proportion of neurons alive, and thus being able to examine the effects on organoids’ electrophysiological firing parameters. An anti-QUIN concentration of 10 *µ*g*/*mL was used based on our previous study of the anti-QUIN’s ability to rescue oligodendrocyte cell lines from QUIN toxicity^7^. QUIN was also produced by the organoids endogenously which were quantified by Gas chromatography–mass spectrometry (GC-MS) in the range 16-30 nM (Supplementary Fig.8).

#### MEA recordings

The organoids from each cell line were plated on a separate multi-well MEA plate, resulting in a total of four plates corresponding to the four cell lines. Within the multi-well MEA, each organoid was placed in a distinct well, where each well was composed of a 3 x 3 grid of titanium nitride electrodes. Each electrode has a diameter of 30 *µ*m with an inter-electrode distance of 200 *µ*m. The electrodes in each well recorded the electrophysiological activity of the organoid in it at varying time intervals. MEA data were recorded at a sampling frequency of 25 kHz using the Multi-Channel Systems data acquisition system. At each sampling time point (represented by a row in Table 2), the media was changed and the recordings were taken before the media change.

#### Increment formation

The data increments were formed based on the sampling timepoints of the captured MEA recordings. Specifically, the pre-processed MEA signal segments obtained at sampling timepoint *i* for a given cell line constituted the *i*-th increment (*I*_*i*_) specific to that cell line (Table 2). For each cell line, *i* = 1 is when the first measurement was recorded and 1 ≤ *i* ≤ 6. Note that the number of increments (the range of *i*) vary for each cell line, but this does not interfere with our modeling because we only compare and model the organoids from the same cell line.

#### MEA Pre-processing

The recorded MEA signals were passed through a third-order high-pass Butterworth filter at 300 Hz to remove low-frequency artifacts^23^. Each recording was segmented into 4-second non-overlapping windows. The 4-second segment length was empirically determined by experimenting with varying segment lengths. Specifically, we conducted a systematic exploration of different window lengths, ranging from 1 second to 20 seconds, considering the typical total length of the signal, approximately 2 minutes. Our experiments took into account various factors, including the duration of events of interest such as spikes or bursts, the computational cost of the algorithm, and the quality of the resulting visualizations. Based on these considerations, we found that a 4-second window length yielded optimal results. MEA signals are often distorted with noise and disturbance which are not well understood in the time domain as it is in the frequency domain. Therefore, we used FFT to represent the signal segments in the frequency domain^8^. The FFT is an efficient implementation of the discrete Fourier transform. Once the FFT of the MEA signal segment was obtained, we computed the Fourier amplitude spectrum of the discrete frequencies. Since the MEA signals are real-valued, the obtained Fourier Transformation is conjugate symmetric. Therefore, we only used the first half of the amplitude spectrum while leaving out the second half. We then reduce the number of features in the frequency domain to 1000 before applying dimensionality reduction.

### Simulated Dataset

#### Trajectory formation

To simulate random non-linear trajectories, we propose the following method. We first generate a line *L* and subsequently transform it to a non-linear trajectory *T* (both *L* and *T* are in D-dimensional space). Specifically, *L* originates from a point *p* and has the gradient *d*. The pseudo-temporal parameter *t* is used to describe points along the line *L*. The vector form of *L* can be represented as shown in Equation 1 where the intercepts (*p*_1_, *p*_2_, …, *p*_*D*_) and the coefficients (*d*_1_, *d*_2_, …, *d*_*D*_) were randomly generated.

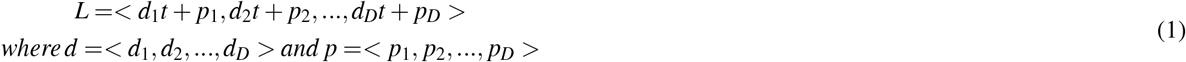

*L* was then transformed into a non-linear trajectory *T* by taking the square of the components in randomly selected dimensions (Equation 2).

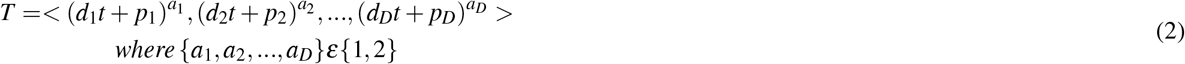

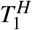 and 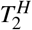 were created from Equation 2 using two different random initializations. In addition to 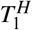 and 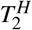, which we assume to be independent of each other, there is the need to create trajectories that exhibit some similarity or show some dependence on each other to verify IL-VIS’s capability in capturing (dis)similarities in trajectory progressions. Therefore, 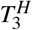 and 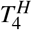 were defined as compositions of 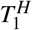 and 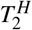 (Equation 3). The similarity factor (*α*) determines how closer the secondary trajectory (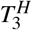 or 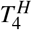) is to its principal trajectory (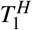or 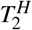).

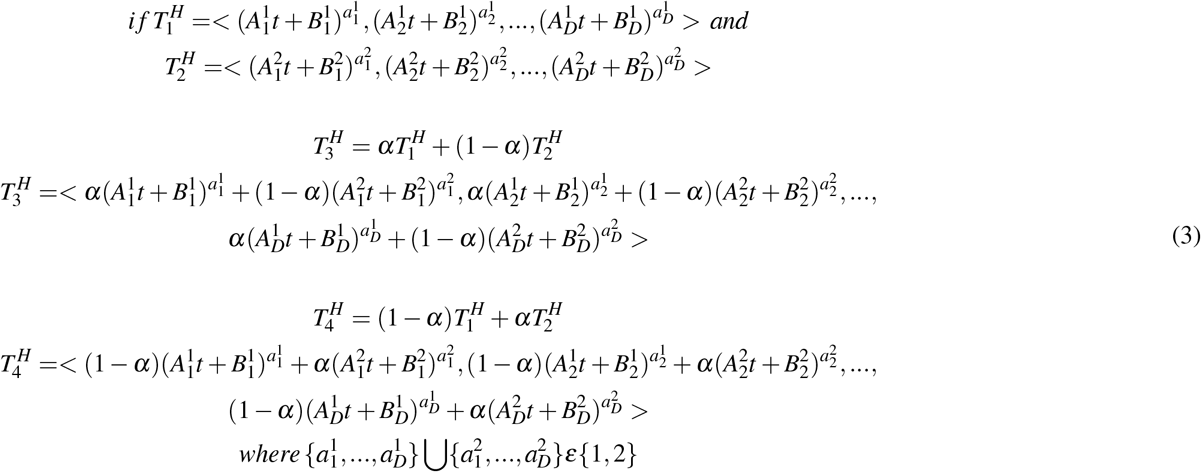

#### Increment formation

In the simulation experiments, the data increments (*I*_1_,…,*I*_*n*_) were formed by generating a set of points along the trajectory 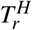 at varying intervals of *t* that are disjoint and increasing. For each increment *I*_*i*_, an interval was defined (Equation 4) based on *i* and the sampling gap size *δ*. Consequently, for each integer *t* value, we created one data point, resulting in a total of (100 *−δ*) data points per increment.

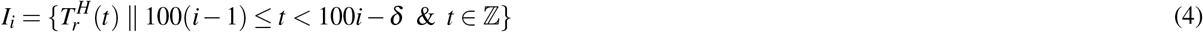

For example, in experiments without a sampling gap (*δ* = 0), 100 data points were generated for each data increment. The *t* value varied from 0 to 99 for the first increment (*i* = 1), and from 100 to 199 for the second increment (*i* = 2) where subsequent increments followed the same pattern. Similarly, in experiments with a sampling gap of size 25, 75 data points were generated for each increment. The *t* value varied from 0 to 74 for the first increment (*i* = 1), from 100 to 174 for the second increment (*i* = 2) and subsequent increments followed the same pattern.

#### Noise addition to the trajectory

We introduce noise for two purposes. First, to approximate random noise arising from biological and non-biological variations in real data, *ϕ*_*i*_(*t*) was added to the data points in each increment *I*_*i*_ (Equation 5). Specifically, for each *D*-dimensional data point, *ϕ*_*i*_(*t*) was added independently to each dimension. *ϕ*_*i*_(*t*) was sampled from a Gaussian distribution with a mean of 0 and a standard deviation (SD) specific to each dimension. To vary the amount of noise in each experiment, *ϕ*_*i*_(*t*) was scaled by the parameter *λ*. *λ* allows for control over the magnitude of the noise added to the data points.

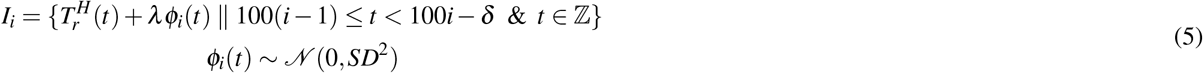

Second, for the experiment with the largest sampling gap (when *δ* = 99), the process described in Section “Increment Formation” would only yield one data point per increment. It is important to note that our objective is not to interpolate values between the sampled points but rather to overcome this limitation and obtain multiple data points per increment. We add noise to the above point generating 100 points per increment. This noise was sampled from a Gaussian distribution with a mean of 0 and an SD of 0.001.

### Incrementally learned dimensionality reduction module

#### Dimensionality reduction algorithm

IL-VIS needs an accurate dimensionality reduction algorithm that caters to incrementally gathered high-dimensional data. This algorithm should be capable of retaining a set of parameters that can be updated over time as new data is incorporated. To this end, we selected our recently proposed parametric algorithm SONG.

The functionality of SONG can be summarised in three primary iterative steps: 1) vector quantization, 2) self-organization, and 3) dimensionality reduction. In vector quantization, the high-dimensional input space *X* = {*x* ∈ *ℜ*^*D*^} is partitioned based on the high-density regions in the input distribution, and each partition is represented by a high-dimensional coding vector, *c* ∈ ℜ^*D*^. In the self-organization step, the set of coding vectors *C* = {*c* ∈ ℜ^*D*^}, are connected based on their proximity to approximate the input topology, effectively parametrizing the input space. An edge between two coding vectors is assigned a higher weight if a dense region of input points lies in between the neighborhoods of the two connected coding vectors, and a lower weight if otherwise. The resulting coding vectors *C* and their adjacency matrix *E* create a weighted graph *G* = {*C, E*}, that could be used to generate a neighborhood distribution. In the dimensionality reduction step, a set of 2D vectors *Y* = {*y* ∈ *ℜ*^2^} which has a bijective correspondence with *C* are defined, i.e., the input *x*_*k*_ ∈ *ℜ*^*D*^ is visualized as *y*_*k*_ ∈ *ℜ*^2^. By minimizing the cross entropy between the neighborhood graph in the input space and the 2D representations, the topology of the input space is then projected onto the 2D space. For more detailed information on SONG refer to work^5^.

Unlike many other dimensionality reduction methods, SONG can retain the information about the data it has seen previously by means of retaining the parameters at time *t* (*θ*_*t*_). *θ*_*t*_ consists of the coding vectors *C*^*t*^, the edges *E*^*t*^, and the corresponding low-dimensional vectors *Y*^*t*^ at time *t* (*θ*_*t*_ = {*C*^*t*^ ∪*E*^*t*^ ∪*Y*^*t*^}). When new data is presented, rather than arbitrarily reinitializing all the relevant visualizations, SONG fine-tunes *θ*_*t*_ to adapt to the new data. This allows SONG to adjust existing visualizations and interpretations to suit the changes coming with the new data, thereby making it more suitable for longitudinal studies.

#### Incremental Training

The dimensionality reduction model is incrementally trained for *n* sessions, one each per sampling time point, where at each session *i* the newly generated data from the current sampling timepoint *i* is incorporated into the existing model. When incrementally training a parametric model using only the newly encountered data, it experiences a phenomenon known as catastrophic forgetting^36^. The model may forget previously learned knowledge as it acquires new knowledge. Therefore, at each session *i*, we perform fine-tuning on the existing parameters *θ*_*i−*1_ using the combined data from all the sampling time points up to and including time point *i*, denoted as 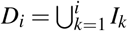. This process allows us to obtain updated parameters *θ*_*i*_, mitigating forgetting and ensuring the generation of reliable visualizations.

At the end of each training session *i*, the updated parameters *θ*_*i*_ are used to transform *D*_*i*_ into 2D representations which are then visualized through a scatter plot.

#### Median tracing

The medians of the visualized data increments are calculated using the 2D coordinates of the representative data points. The calculated medians are then traced chronologically to observe the emergence of a trajectory in the 2D space. The median was used instead of the mean as medians are less impacted by possible outliers.

## Euclidean Distance Metric

A Euclidean distance-based metric was used to quantify the (dis)similarity of progressions of any two trajectories. The metric calculates the average Euclidean distance between the medians of the corresponding increments in the considered trajectories. Equation 6 calculates the distance between two 2D trajectories A and B, each with *n* increments. 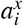 and 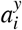 correspond to the medians of the dimensions *x* and *y* in the *i*^*th*^ increment of trajectory A. 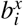 and 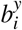 correspond to the medians of the dimensions *x* and *y* in the *i*^*th*^ increment of trajectory B.

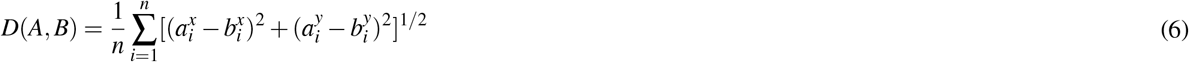

### Spike detection

MEA data were also analyzed using Neuroexplorer v5 software tool^37^. The reference channel was subtracted, a band pass filter (Butterworth) from 300 to 1000 Hz was applied and spikes were detected using the threshold crossing algorithm. The threshold was calculated as -4.0*Median Sigma (MS), whereby MS is equal to the median of the signal absolute values.

## Code Availability

All code related to simulation trajectory formation, MEA data pre-processing and the proposed incremental pipeline could be found in the following repository https://github.com/TamashaM/IncrementalFramework.

## Data Availability

Experimental data supporting the findings of this paper are available from L.O. upon request.

## Supplementary information

The supplementary material related to this manuscript can be found in the additional document attached to the submission.

## Acknowledgments

The authors acknowledge Dementia Australia Research Foundation and Yulgilbar Alzheimer’s Research Program for funding.

T.M. acknowledges Melbourne Graduate Research Scholarship and GCI Women in STEM Student Award support scheme.

D.S. and S.H. acknowledge Australian Research Council grant DP210101135. L.O. is supported by a National Health and Medical Research Council (NHMRC) of Australia Boosting Dementia Research Leadership Fellowship (APP1135720). V.G. acknowledges Australian Research Council Discovery Early Career Researcher Award DE180100775. M.D.L., G.S., and B.B. acknowledge funding of the Peter Duncan Neurosciences Research Unit at St. Vincent’s Centre for Applied Medical Research and funding from a Perpetual IMPACT grant. M.D.L., G.S., and B.B. acknowledge the contribution of Prof. Daniel Christ and Dr. Peter Schofield toward the expression and purification of the IgG1 antibody used in this study. The authors wish to thank Prof. Perminder S. Sachdev (Centre for Healthy Brain Aging [CHeBA], University of New South Wales, Australia) for provision of the fibroblast lines that were reprogrammed to iPSCs.

The authors would like to thank Nisal Ranasinghe, Deshani Geethika, Bishakhdatta Gayen, Peter Savas, Maneesha Perera, Rashindrie Perera, Hai-Hang Wu, Jayanie Bogahawatte and Nadarasar Bahavan for proofreading.

## Author contributions statement

T.M., D.S., V.G., M.E., M.D.L., G.S., B.B., C.J., L.O., and S.H. designed research; T.M., R.B. and S.C. performed research; T.M., D.S., V.G., and S.H. analyzed data (ML); M.E., M.D.L., B.H., S.C. and L.O. analyzed data; T.M., D.S., M.D.L., G.S., C.M, G.G. and B.B., contributed new reagents/analytic tools; M.E., and R.B., cultured cells; T.M., D.S., V.G., M.D.L., G.S., R.H., S.C., C.M, G.G., B.B, L.O., and S.H wrote the paper.

