## Supplementary materials V2 for "Visualization of Incrementally Learned Projection Trajectories for Longitudinal Data"

### Supplementary Material

May 31, 2024

#### Human brain organoid production

Human brain organoids [2] produced from iPSCs based on the protocol by Salick et al. [3] were cultured for up to 9 months. Briefly, iPSC colonies were passaged using PBS-EDTA as near single cell suspension, plated onto matrigel (Corning #354277) coated 60 mm cell culture dishes (5000 cells per dish) and maintained in TeSR-E8 media (StemCell Technologies, #5990) for five days in hypoxic conditions (3% $O_2$, 5% $CO_2$, 37°C). Five day old iPSC colonies were collected via PBS-EDTA (Thermo Fisher Scientific #14190-144, #15575-038), dissociated into a single cell suspension, and seeded at 1000 cells per well into U-bottom 96 well plates (Corning #7007) in iPS-Brew (Miltenyi Biotec, #130-107-086) containing 10 $\mu$M ROCK inhibitor (Y-27632, Calbiochem #688000), followed by brief centrifugation (2 min at 400 G) to focus the cell suspensions. Partial media changes were performed every 48h. Embryoid body media was used for days 2 to 11 (DMEM/F12 (Thermo Fisher Scientific #12500-096) supplemented with 1% NEAA (Thermo Fisher Scientific #11140-050), 1% Glutamax (Thermo Fisher Scientific #35050-061), 1% B27 (Thermo Fisher Scientific #17504-001), 1% N2 (Thermo Fisher Scientific #17502-001)) and contained 10 $\mu$M SB431542 (Stemcell Technologies #72234), 0.1 $\mu$M LDN193189 (MedChemExpress #HY-12071) and 1 $\mu$M IWP-2 (MedChemExpress #HY-13912) until day 8, and 20 ng/$\mu$l FGF2 (Stemcell Technologies #130-104-923) and EGF (Miltenyi Biotec #130097750) until day 11. Neural induction was triggered by phasing in neural media (Brainphys (Stemcell Technologies #05790) containing 1% NEAA, 1% Glutamax, 2% B27, 1% N2 until day 24) with partial media changes every 48h until day 24. Spheroids were then transferred to 24 well plates to accommodate the larger media volume needs, maintained in normoxic conditions (20% $O_2$, 5% $CO_2$, 37°C), and matured through the addition of 5 ng/ml BDNF (Miltenyi Biotec #130-096-286) and 10 ng/ml GDNF (Miltenyi Biotec #130-098-449) from day 26, and 10 ng/ml NRG1 (Peprotech #AF-100-03) and 20 ng/ml NT3 (Miltenyi Biotec #130-093-973) from day 34, after which partial media changes were performed every 3 to 4 days until the end of the experiment.

#### Anti-Quinolinic Acid antibody production and purification

The monoclonal anti-QUIN (clone 7AR, mouse anti-human, isotype IgG1) was developed by Millipore, and subsequently patented by B.J.B. and G.J.G (PCT/IB2013/055902 and WO/2015/008111). Briefly, hybridoma clones producing the antibody were grown in Hybridoma-SFM serum-free medium and scaled up to a volume of 500 mL. After low-speed centrifugation (4,000×g at 4°C for 10 min) and sterile filtration (with a 0.22 $\mu$m filter), the antibodies were purified using a column packed with Protein G Sepharose 4 fast flow beads (GE Healthcare). Elution was carried out with 0.1 M glycine (pH 2.7), and the eluate was neutralized with 1 M Tris-HCl (pH 8.5), after which buffer exchange to PBS at pH 7.4 was carried out. Purified antibodies were quantified using spectrophotometer by measuring the absorbance at 280 nm. Affinity of interactions between biotinylated antibody and purified protein was measured by BLI (BLItz, ForteBio), at room temperature.

#### Separability of the trajectories with a larger dataset

In this section, we explore the performance of IL-VIS on larger datasets. Since increasing the number of samples in real data is infeasible due to the considerable time and cost associated, we conduct an additional experiment in which we increase the number of simulated trajectories. Specifically, the

Table 1: Distance between simulated trajectories when $\alpha = 0.8$

| | $T_1^L$ | $T_2^L$ | $T_3^L$ | $T_4^L$ |
|---|---|---|---|---|
| $T_1^L$ | x | 2.72 | 0.30 | 2.62 |
| $T_2^L$ | x | x | 2.77 | 0.34 |
| $T_3^L$ | x | x | x | 2.64 |
| $T_4^L$ | x | x | x | x |

Distance between the trajectory pairs after the final session of training calculated using Equation 6 in Methods. $T_3^L$ is closer to $T_1^L$ than to $T_2^L$, and $T_4^L$ closer to $T_2^L$ than to $T_1^L$. $T_1^L$ is the farthest from $T_2^L$

number of simulated trajectories was incrementally raised to a maximum of 20 trajectories and in each experiment, the trajectories were designed to originate at the same location. The results are shown in Fig. 7. Up until the number of trajectories is 10, we can see an obvious separation between their progression. Although they overlap at the initiation points, this is expected as they originate at the same position and are similar at the beginning. As the number of trajectories increases, it becomes increasingly difficult to show clear separation between the trajectories. However, it is important to note that the distinguishability of trajectories is not solely determined by their quantity but is also shaped by factors such as their similarity and length. We recognize that managing a higher volume of trajectories poses a significant challenge, and we believe that this challenge is not unique to our method alone.

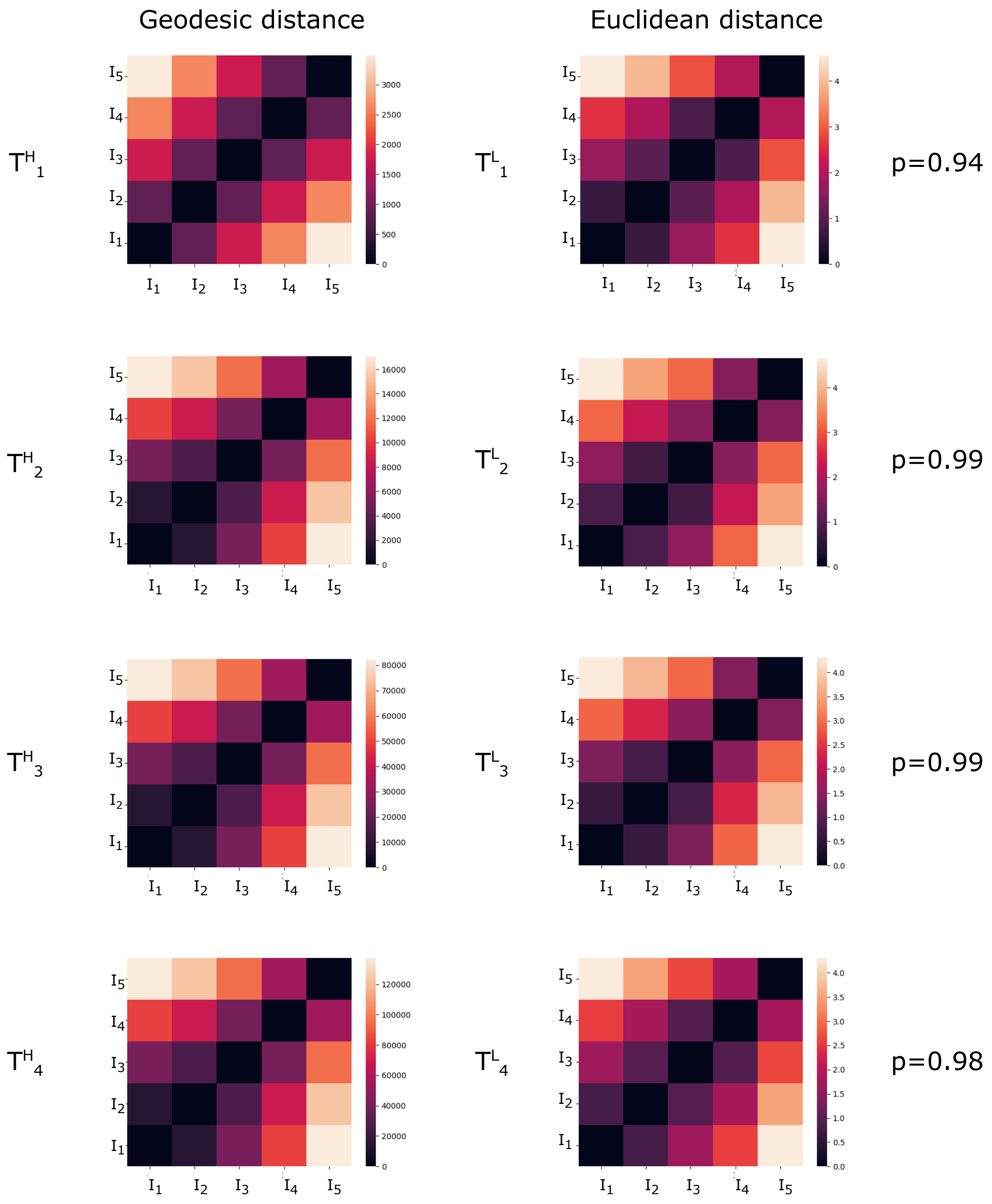


Figure 1: (A) Heatmap of the pairwise geodesic distance between increments calculated using the medians of the 100-dimensional coordinates in each increment in $T_r^H$ confirmed that the $i^{th}$ increment $(I_i)$ is placed away from its previous increments $(I_1, ..., I_{i-1})$ in the high-dimensional space. (B) Heatmap of the pairwise Euclidean distance between increments in $T_r^L$ calculated using the median of the 2D coordinates in each increment showed that the $I_i$ is visualized away from its previous increments $(I_1, ..., I_{i-1})$ in the low-dimensional space similarly to $T_r^H$. We further verified the manifold preservation quality in the SONG embedding by higher Pearson correlation coefficient values (p) between the pairwise Euclidean distances in the visualization space and their corresponding pairwise geodesic distances in the high-dimensional space.

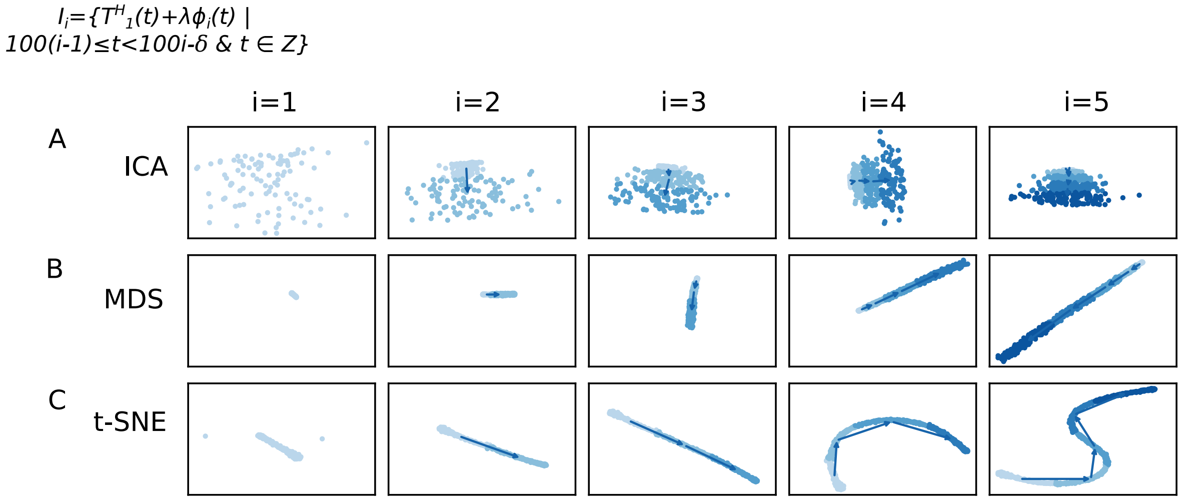


Figure 2: 2D visualizations obtained by the pipeline when modeling $T_1^H$ using (A) ICA, (B) MDS, and (C) t-SNE as the dimensionality reduction technique. $T_1^H$ was defined on a pseudo-temporal parameter $t$ (Equation 2 in Methods). $\phi_i(t)$ is noise sampled from a Gaussian distribution (Methods). $\lambda$ is the percentage of noise added which is 20% for this experiment. Five increments of data points are formed ($n = 5$) for each trajectory with no sampling gaps ($\delta = 0$) in between thus each increment contains 100 data points (see Methods). $N$ corresponds to the number of data points used in each visualization. Each row contains the five visualizations generated at the five sessions where a new data increment is introduced at each session. At session $i$, all the increments thus far (i.e., $D_i = \bigcup_{k=1}^{i} I_k$) are visualized. In visualizations with multiple increments, different increments are displayed in varying intensities of the same color. The stronger the intensity, the newer the increment. By tracing the medians of the subsequent increments chronologically, the emergence of $T_1^L$ which is a 2D representation of $T_1^H$ is observed. (A and B) Both ICA and MDS failed to generate progressive visualizations where the orientation of the formed trajectories changes over the sessions. Further, they both failed to preserve the non-linearity of $T_1^H$ (C) t-SNE preserved the non-linearity of $T_1^H$ but failed to preserve the orientation of the trajectory over the incremental sessions.

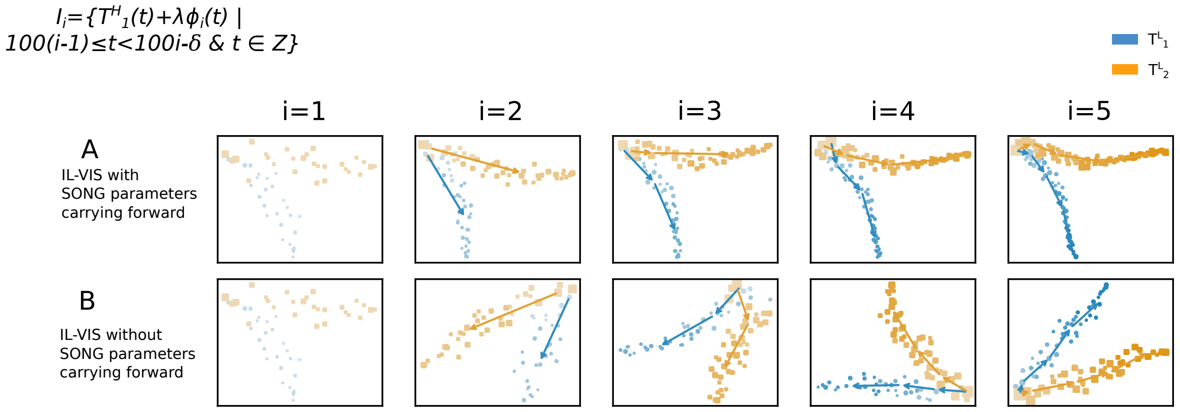


Figure 3: 2D visualizations obtained by IL-VIS for two randomly generated trajectories $T_1^H$ and $T_2^H$ (A) when parameters of previous iterations are carried forward and (B) when parameters of previous iterations are not carried forward. $T_1^H$ and $T_2^H$ and was defined on a pseudo-temporal parameter $t$ (Equation 2 in Methods). $\phi_i(t)$ is noise sampled from a Gaussian distribution (Methods). $\lambda$ is the percentage of noise added which is 0% for this experiment. Five increments of data points are formed ($n = 5$) for each trajectory with no sampling gaps ($\delta = 0$) in between thus each increment contains 100 data points (see Methods). $N$ corresponds to the total number of data points used in each visualization. Each experiment contains five visualizations generated at the five sessions. At each session $i$, the $i^{th}$ data increment of all trajectories that are modeled together is introduced to the pipeline. Each trajectory is shown in a different color. In visualizations with multiple increments, different increments are displayed in varying intensities of the same color. The stronger the intensity, the newer the increment. By tracing the medians of the subsequent increments chronologically, the emergence of $T_r^L$ which is a 2D representation of $T_r^H$ is observed. (A) When parameters are carried forward which is the default implementation, IL-VIS generates evolving visualizations preserving the same orientation over the incremental sessions. (B) When parameters are not carried forward, i.e., when SONG is reset at each session IL-VIS cannot preserve the orientation of the trajectories and would thus move around the visualization space challenging the mapping of the visualizations generated at different sessions.

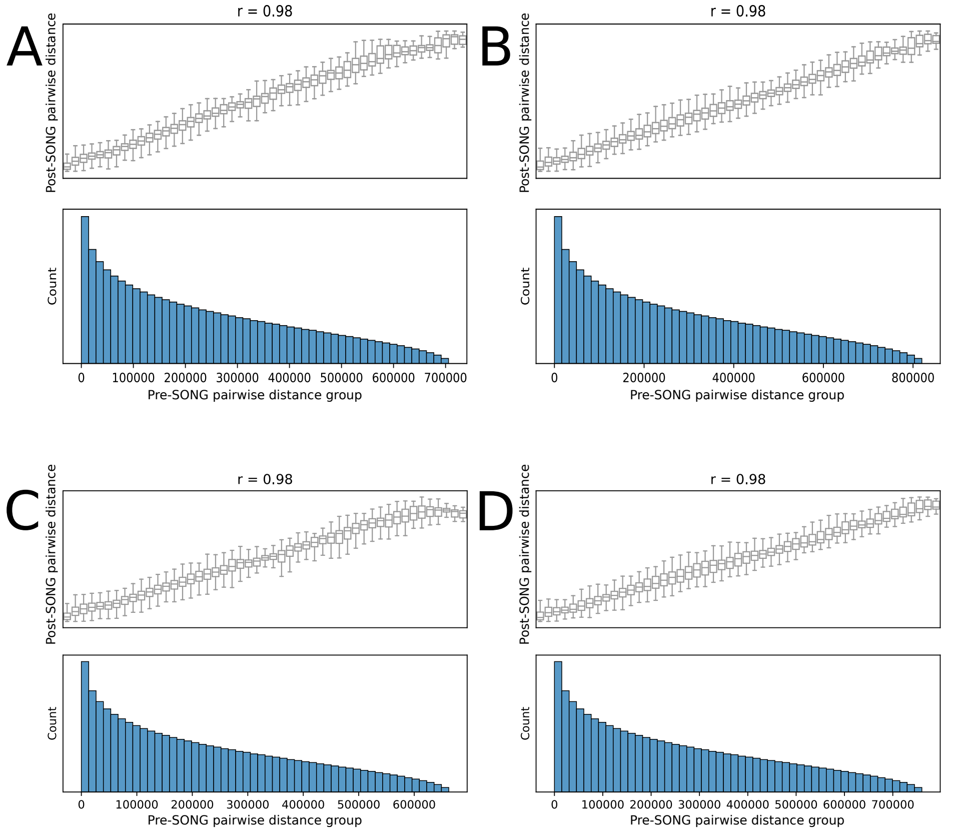


Figure 4: We adapted the experiment described in [1] to evaluate the pairwise distance preservation in low-dimensional SONG embeddings. Subfigures correspond to the pairwise distance preservation in the low-dimensional space for (A) $T_1^H$ (B) $T_2^H$, (C) $T_3^H$ and (D) $T_4^H$ at the fifth session ($i = 5$). Each box plot represents the distribution of the low-dimensional pairwise distances plotted against their corresponding high-dimensional pairwise distance bin. Bins are created using 50 equal width bins over the pairwise distances in the original 100-dimensional space. The frequency of original pairwise distances in each bin is shown below each boxplot. The computed Pearson correlation coefficient over the pairs of pairwise distances is marked above each graph. We observed that SONG has been successful in preserving the small to large-scale high-dimensional distances in the low-dimensional space.

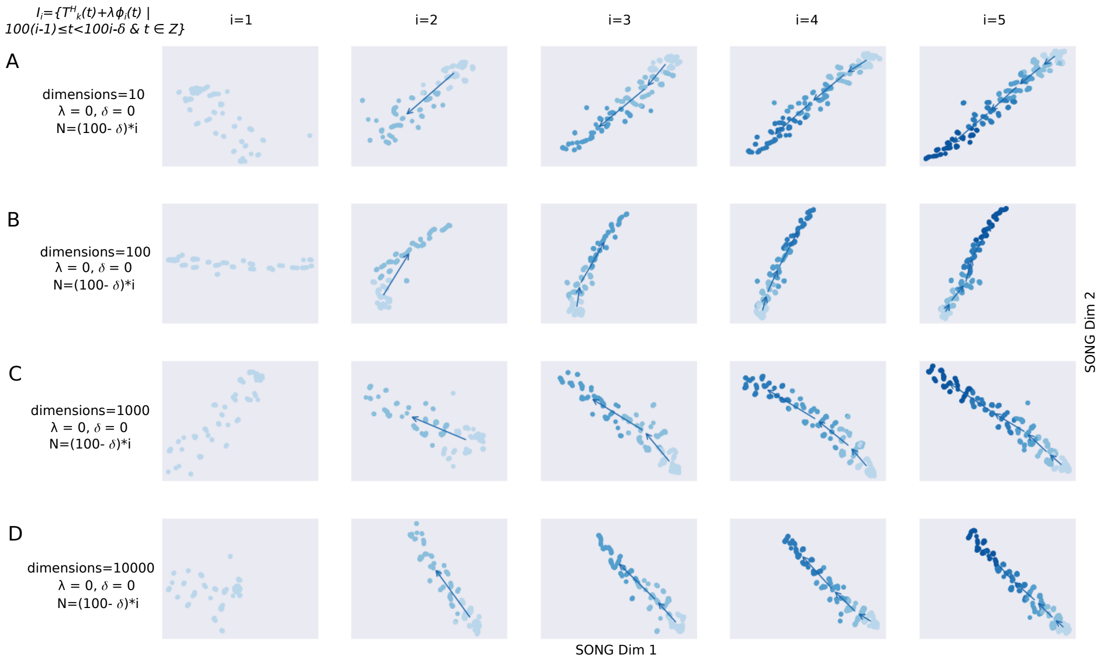


Figure 5: 2D visualizations obtained by IL-VIS showed its capability in catering for various number dimensions (A) 10 dimensions, (B) 100 dimensions, (C) 1000 dimensions, and (D) 10000 dimensions. $\phi_i(t)$ is noise sampled from a Gaussian distribution (Methods). $\lambda$ is the percentage of noise added which is 0% for this experiment. Five increments of data points are formed ($n = 5$) for each trajectory with no sampling gaps ($\delta = 0$) in between thus each increment contains 100 data points (Methods). $N$ corresponds to the number of data points used in each visualization. Each row contains the five visualizations generated at the five sessions where a new data increment is introduced at each session. At session $i$, all the increments thus far (i.e., $D_i = \bigcup_{k=1}^{i} I_k$) are visualized. In visualizations with multiple increments, different increments are displayed in varying intensities of the same color. The stronger the intensity, the newer the increment. By tracing the medians of the subsequent increments chronologically, the emergence of $T_k^L$ which is a 2D representation of $T_k^H$ is observed. The visualizations confirmed the IL-VIS's capability to produce evolving visualizations while capturing the gradually increasing and non-linear trend of simulated trajectories even with a higher number of dimensions.

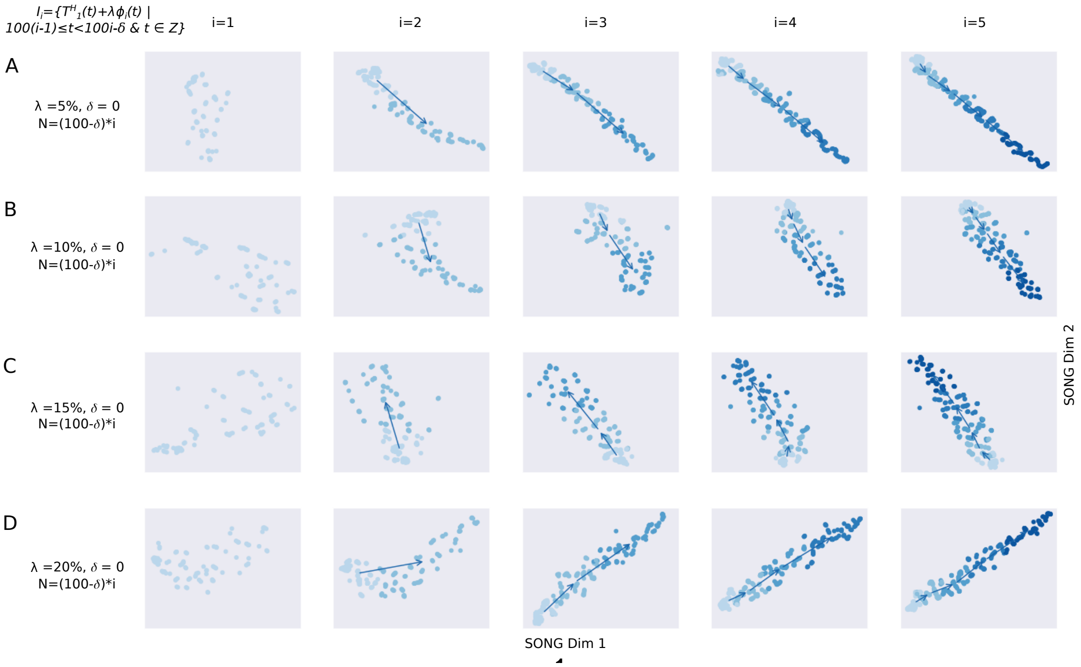


Figure 6: 2D visualizations obtained by IL-VIS showed its robustness to various degrees of noise. (A) 5%, (B) 10%, (C) 15%, and (D) 20%. $\phi_i(t)$ is noise sampled from a Gaussian distribution (Methods). $\lambda$ determines the percentage of noise added which is varied from 5-20% in (A)-(D). Five increments of data points are formed ($n = 5$) for each trajectory with no sampling gaps ($\delta = 0$) in between thus each increment contains 100 data points (Methods). $N$ corresponds to the number of data points used in each visualization. Each row contains the five visualizations generated at the five sessions where a new data increment is introduced at each session. At session $i$, all the increments thus far (i.e., $D_i = \bigcup_{k=1}^{i} Ikj$) are visualized. In visualizations with multiple increments, different increments are displayed in varying intensities of the same color. The stronger the intensity, the newer the increment. By tracing the medians of the subsequent increments chronologically, the emergence of $T_1^L$ which is a 2D representation of $T_1^H$ is observed. Despite the presence of this added noise, $T_1^L$ successfully preserves the trend and non-linearity of $T_1^H$. However, the smoothness of $T_1^L$ decreased as the noise percentage increased.

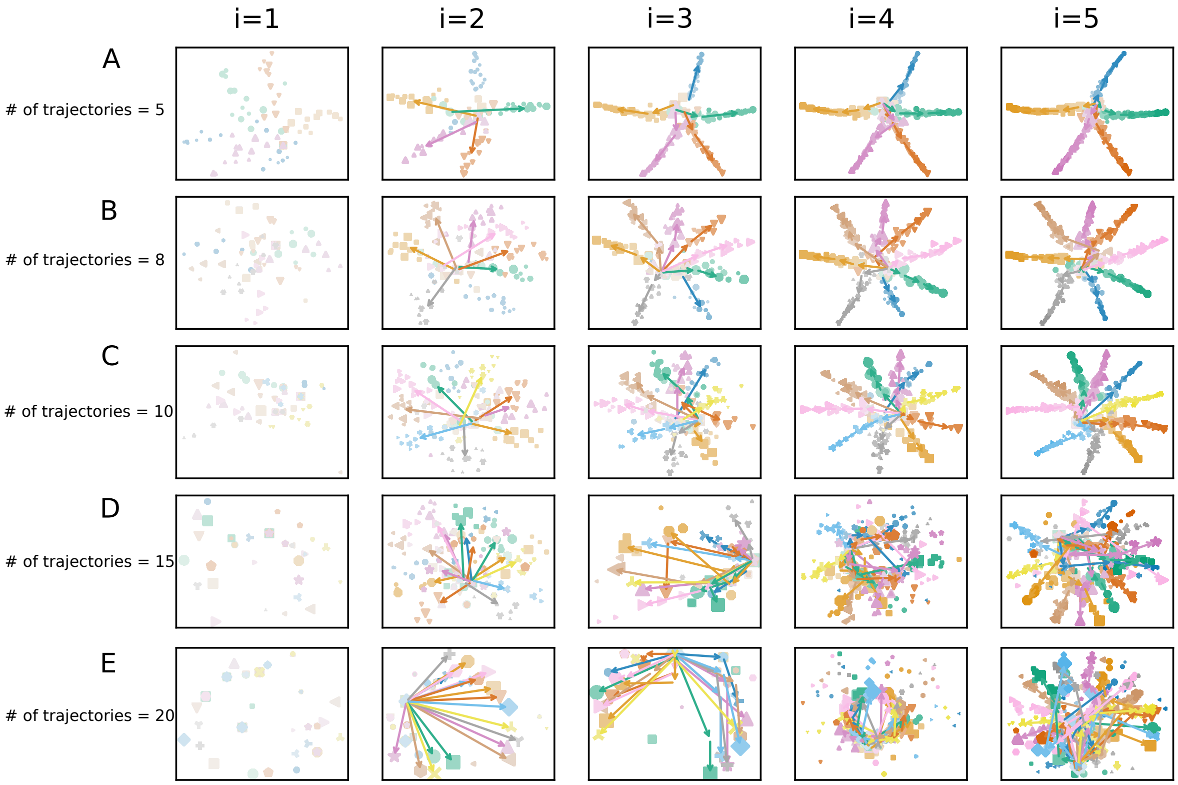


Figure 7: ]2D visualizations obtained by IL-VIS when the number of trajectories ($R$) in each experiment was (A) 5, (B) 8, (C) 10, (D) 15 and (E) 20. All the trajectories are 100-dimensional non-linear trajectories that were randomly generated. Five increments of data points are formed ($n = 5$) for each trajectory with no sampling gaps. The percentage of noise ($\lambda$) added was also 0% for this experiment. Each experiment contains five visualizations generated at the five sessions. At each session $i$, the $i^{th}$ data increment of all trajectories that are modeled together is introduced to the pipeline. Each trajectory is shown in a different color. In visualizations with multiple increments, different increments are displayed in varying intensities of the same color. The stronger the intensity, the newer the increment. By tracing the medians of the subsequent increments chronologically, the emergence of $T_r^L$ which is a 2D representation of $T_r^H$ is observed.

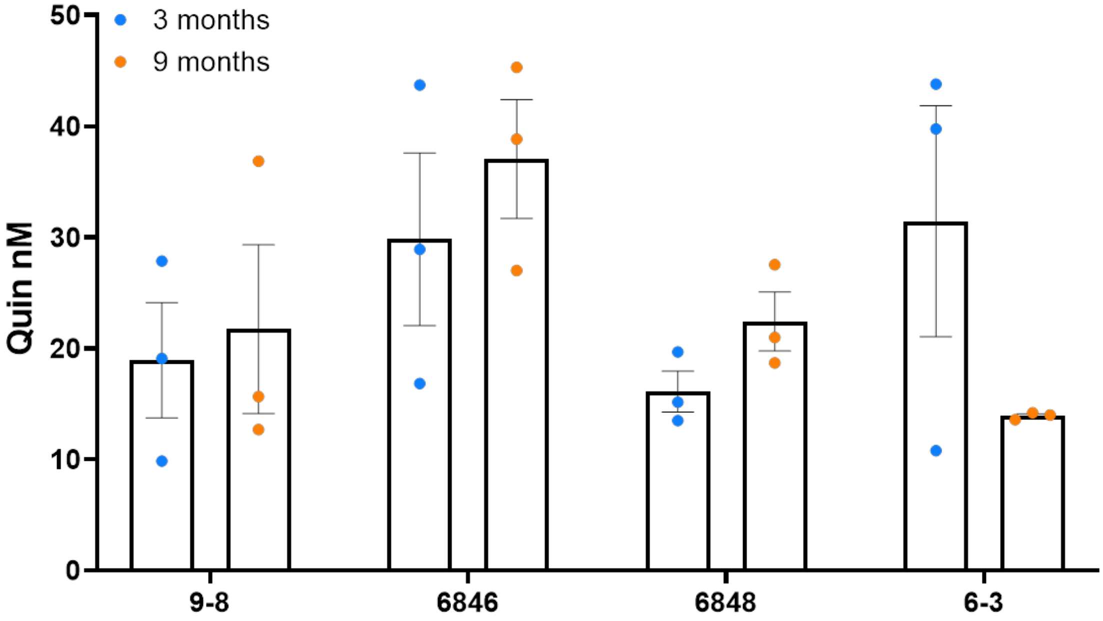


Figure 8: QUIN was produced by the organoids endogenously and released into the media, whereby it was quantified by GC-MS at 16-30 nM

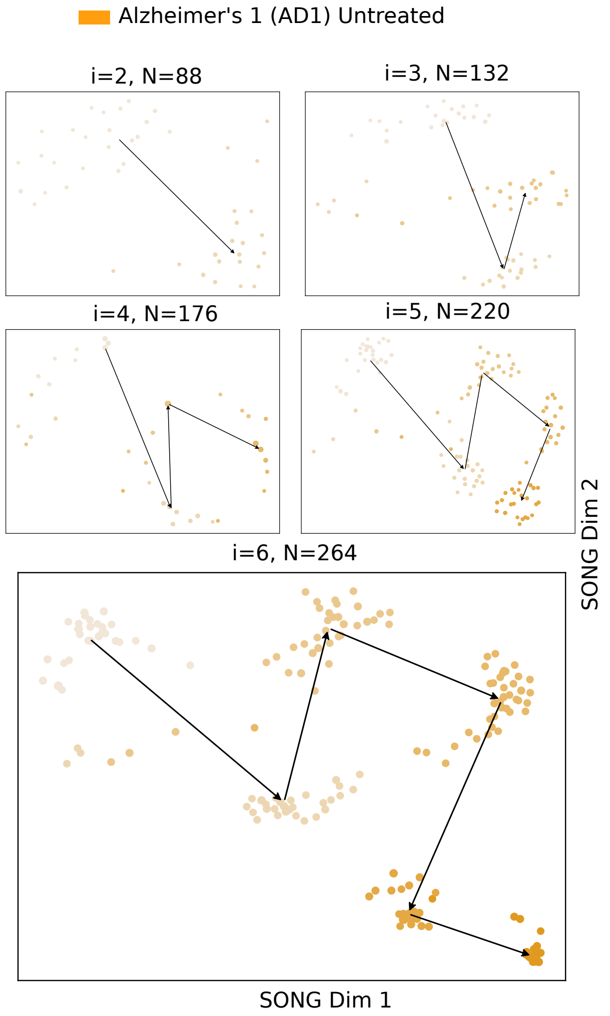


Figure 9: 2D visualizations obtained by IL-VIS for the untreated for Untreated AD1 organoid when modeled independently. $I_1$ to $I_6$ are the six increments corresponding to the six different timepoints at which the recordings were taken (Table 2 in Methods). At session $i$, all the increments thus far (i.e., $D_i = \bigcup_{k=1}^{i} I_k$) are used for training the model. N corresponds to the number of data points used in each visualization where each point corresponds to a compact representation of a 4-second recording obtained from the 9 electrodes in the respective MEA plate. The 2D visualizations generated from $2^{nd}$ session ($i = 2$) onwards are shown in the sub-figures. The color intensity represents the organoid's maturity. In earlier increments, the color is lighter while in later increments the color is darker. The medians of the subsequent increments are traced chronologically to observe the trajectory followed by the organoid's electrophysiological properties.

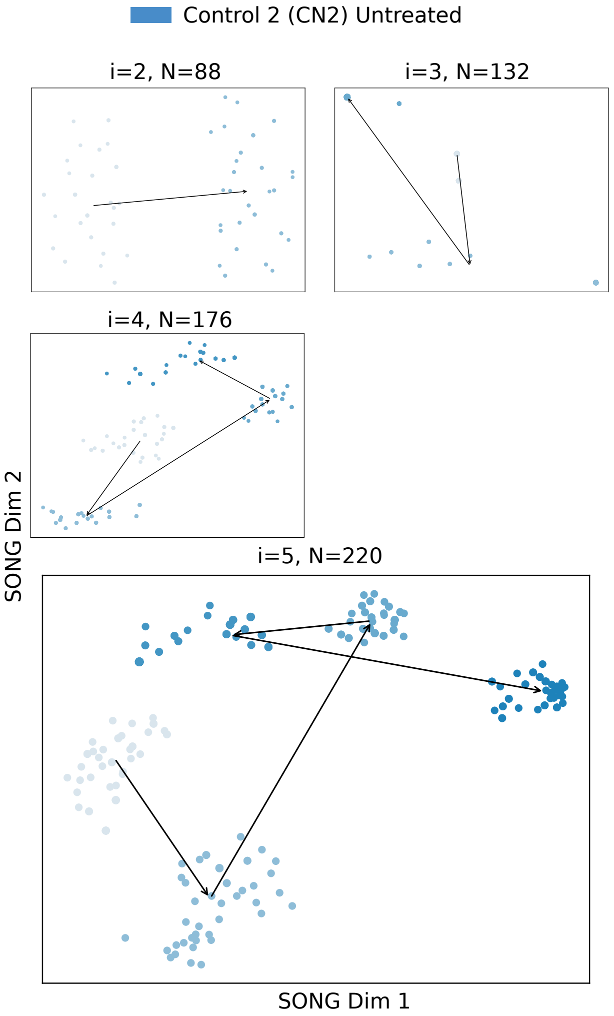


Figure 10: 2D visualizations obtained by IL-VIS for the untreated for untreated CN2 organoid when modeled independently. $I_1$ to $I_5$ are the five increments corresponding to the five different timepoints at which the recordings were taken (Table 2 in Methods). At session $i$, all the increments thus far (i.e., $D_i = \bigcup_{k=1}^{i} I_k$) are used for training the model. N corresponds to the number of data points used in each visualization where each point corresponds to a compact representation of a 4-second recording obtained from the 9 electrodes in the respective MEA plate. The 2D visualizations generated from $2^{nd}$ session ($i = 2$) onwards are shown in the subfigures. The color intensity represents the organoid's maturity. In earlier increments, the color is lighter while in later increments the color is darker. The medians of the subsequent increments are traced chronologically to observe the trajectory followed by the organoid's electrophysiological properties.

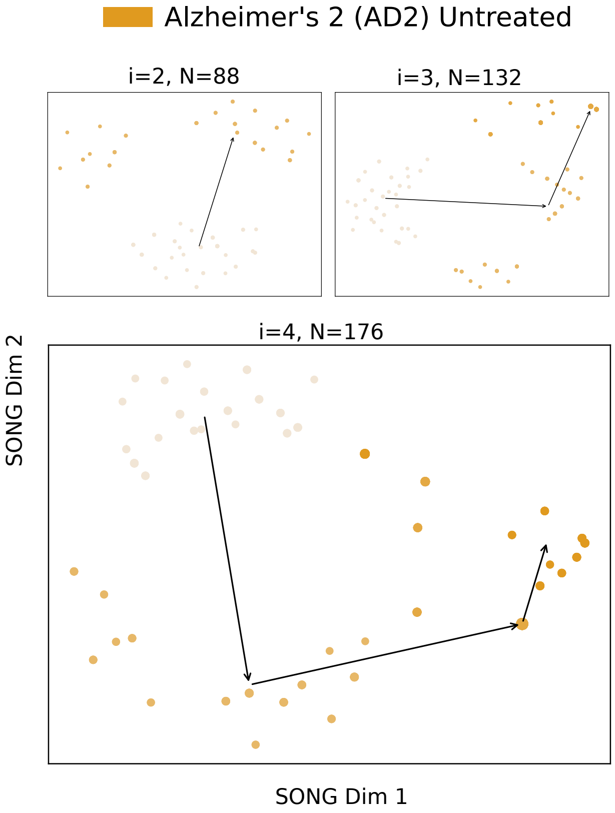


Figure 11: 2D visualizations obtained by IL-VIS for the untreated AD2 organoid when modeled independently. $I_1$ to $I_4$ are the four increments corresponding to the four different time points at which the recordings were taken (Table 2 in Methods). At session $i$, all the increments thus far (i.e., $D_i = \bigcup_{k=1}^{i} I_k$) are used for training the model. N corresponds to the number of data points used in each visualization where each point corresponds to a compact representation of a 4-second recording obtained from the 9 electrodes in the respective MEA plate. The 2D visualizations generated from the $2^{nd}$ session ($i = 2$) onwards are shown in the sub-figures. The color intensity represents the organoid's maturity. In earlier increments, the color is lighter while in later increments the color is darker. The medians of the subsequent increments are traced chronologically to observe the trajectory followed by the organoid's electrophysiological properties.

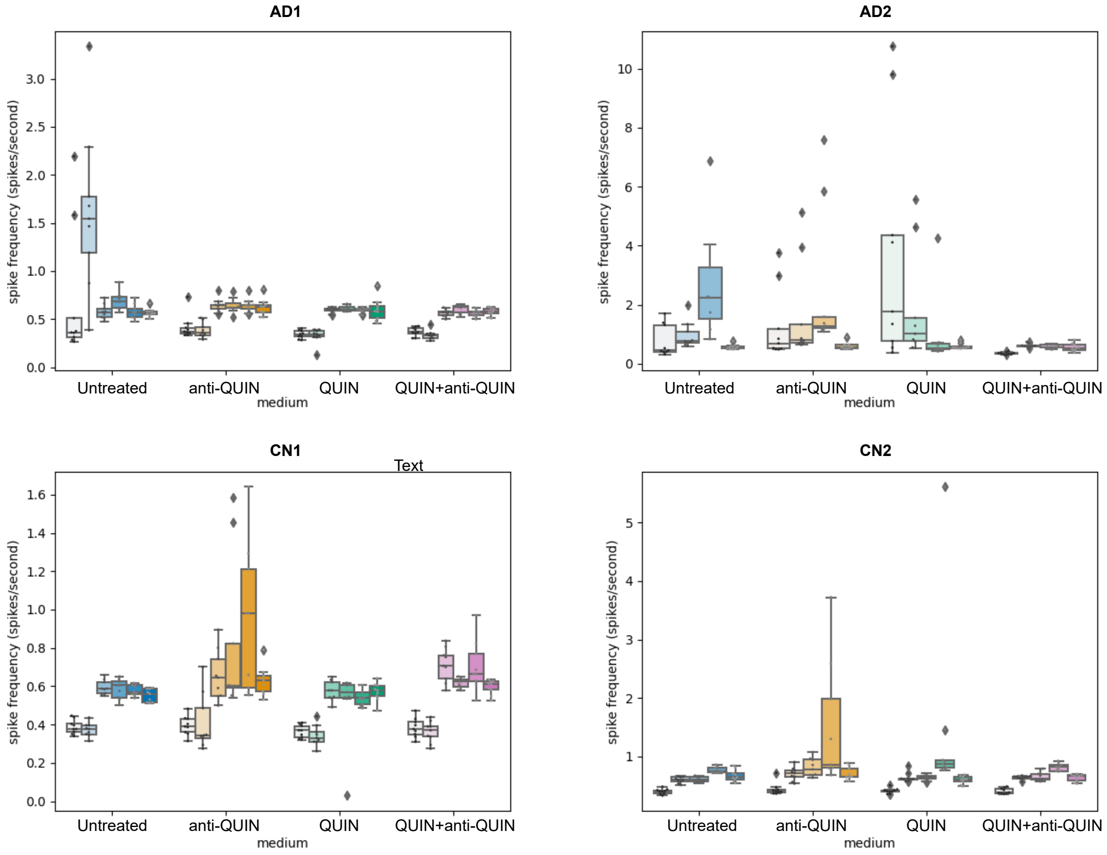


Figure 12: Spiking activity of AD1, AD2, CN1, and CN2 cell lines. The different colors correspond to the different treatments. The boxplot shows how the recorded spike frequency of each organoid in each sampling time point varies in the 9 different electrodes of the corresponding well. The box plots of the same color correspond to the different time points of the same treatment. The color intensity represents the organoid's maturity. At earlier increments, the color is lighter while in later increments the color is darkened. The center line, box, and whiskers correspond to the median, interquartile range (IQR), and minimum–maximum range respectively. Whiskers are extending up to 1.5 times the IQR. Points outside this range are identified as outliers.
